# Analysis of the dynamics of a complex, multipathway reaction: Insulin dimer dissociation

**DOI:** 10.1101/2024.10.08.617297

**Authors:** Kwanghoon Jeong, Spencer C. Guo, Sammy Allaw, Aaron R. Dinner

## Abstract

The protein hormone insulin forms a homodimer that must dissociate to bind to its receptor. Understanding the kinetics and mechanism of dissociation is essential for rational design of therapeutic analogs. In addition to its physiological importance, this dissociation process serves as a paradigm for coupled (un)folding and (un)binding. Based on previous free energy simulations, insulin dissociation is thought to involve multiple pathways with comparable free energy barriers. Here, we analyze the mechanism of insulin dimer dissociation using a recently developed computational framework for estimating kinetic statistics from short-trajectory data. These statistics indicate that the likelihood of dissociation (the committor) closely tracks the decrease in the number of (native and nonnative) intermonomer contacts and the increase in the number of water contacts at the dimer interface; the transition state with equal likelihood of association and dissociation corresponds to an encounter complex with relatively few native contacts and many nonnative contacts. We identify four pathways out of the dimer state and quantify their contributions to the rate, as well as their exchange, by computing reactive fluxes. We show that both the pathways and their extents of exchange can be understood in terms of rotations around three axes of the dimer structure. Our results provide insights into the kinetics of insulin analogues and, more generally, how to characterize complex, multipathway processes.

## Introduction

The kinetics of molecular association and dissociation govern many processes in chemistry and biology ranging from reactions of a few atoms to changes in cell state. The most basic theories of association treat molecules as spheres undergoing Brownian motion;^1^ these theories remain relevant today because they can be extended to account for molecular structure by factoring the rate into terms for reaching an encounter complex and for dynamics within that complex.^1–4^ Despite the success of this approach, it remains difficult to apply when the dynamics in the encounter complex involve significant intramolecular rearrangement, as is often the case for macromolecules such as proteins. In this case, simulations generally cannot directly bridge the separation of timescales between the fastest motions of the molecules (typically bond fluctuations) and the dynamics of interest. A common approach to estimating kinetics when such a separation in timescales exists is to identify a low-dimensional representation and apply transition state theory (TST) and its elaborations that account implicitly for dynamics in the remaining degrees of freedom,^5,6^ but whether the assumptions underlying this approach are valid generally remains unclear (e.g., see Ref. 7 for discussion). Even if they are, this strategy provides limited microscopic insight into the dynamics of the degrees of freedom beyond the reaction coordinates.

These issues are exemplified by studies of the association and dissociation of the monomers in the insulin homodimer. Each monomer consists of two polypeptide chains that are linked by disulfide bridges: a 21-amino acid A-chain and a 30-amino acid B-chain. In the dimer (Figure 1), the secondary structure of the A-chain consists of two *α*-helices (A1–A9 and A13– A20); that of the B-chain consists of an *α*-helix (B9–B19), followed by a *β*-turn (B20–B23), and a *β*-strand (B24–B26). At the dimer interface, the B-chain *α*-helices from the two monomers pack against each other, and the B24–B26 segments form an anti-parallel *β*-sheet. Both experiments^8,9^ and simulations^10–12^ indicate that the A-chain N-terminal helix (A1–A9), the B-chain N-terminal strand (B1–B7), and the B-chain C-terminal residues (B24–B30) become disordered when mutations and/or low pH are used to stabilize the monomer relative to the dimer.

**Figure 1.**
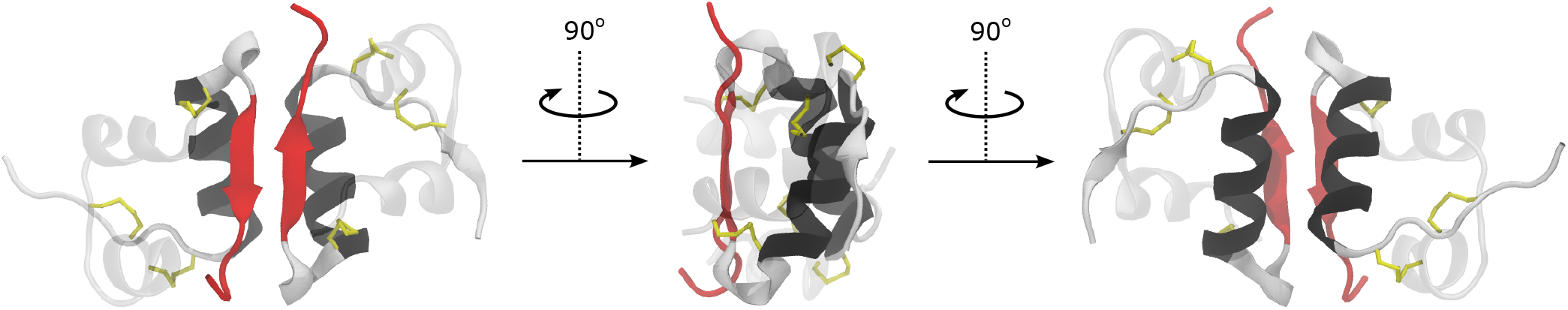
Experimental insulin dimer structure (PDB ID 3W7Y ^13^) from three different points of view. The A-chains are translucent, while the B-chains are opaque. The interfacial *β*-strand and B-chain C-terminal residues (B24–30) are red, and the interfacial *α*-helices (B9–19) are black. Cysteine disulfide bonds are shown in yellow. We use VMD ^14^ to visualize molecular structures throughout the paper.

Dinner and co-workers previously showed that the potential of mean force (PMF) as a function of collective variables (CVs) characterizing the contacts between the interfacial *α*-helices and *β*-strands supports a diversity of dissociation pathways ranging from ones in which the *α*-helices separate first to ones in which the *β*-strands separate first.^15^ Along the limiting path in which the *α*-helices separate first, the *α*-helices twist apart, and water penetrates the hydrophobic core before the *β*-strands slide apart, detach from the core, and become disordered. Along the alternative limiting path, the *β*-strands twist apart first while remaining attached to the core, and the monomers separate largely in their dimeric structures. The paths were estimated to have comparable free energy barriers of around 14 kcal/mol in the dissociation direction and 1– 2 kcal/mol in the association direction. These simulations provide a unified perspective of diverse dynamics in earlier simulations.^11,16–19^

In particular, Bagchi and co-workers inferred a path in which the interfacial *α*-helices separate first from the PMF as a function of the distance between the centers of mass of the monomers and the number of contacts between them.^17^ Based on this two-dimensional PMF and its minimum free-energy path, they subsequently estimated the dissociation rate using a variety of theories; with corrections to the TST rate, they obtained a value of 0.4 *μ*s^−1^.^7^ They identified the transition to the first metastable intermediate with twisted interfacial *α*-helices and a largely intact interfacial *β*-sheet^19^ to be the rate limiting step in TST. The existence of this intermediate is consistent with a twisted dimeric state found in a Markov state model (MSM), though the MSM predicts longer timescales of transition to and from it (*∼*20 *μ*s).^20^ These timescales estimated for the initial events in dissociation are on the order of those detected by temperature-jump two-dimensional infrared (2D IR) spectroscopy of the amide I vibrational modes, which suggestst that monomer disordering is in the 5–250 *μ*s time range and dissociation is in the 250– 1000 *μ*s time range.^21^ On the other hand, time-resolved X-ray scattering measurements suggest dissociation intermediates appear within 1 *μ*s but cannot resolve their structural features.^22^

Given the heterogeneity of possible paths dissociation can take, it is interesting that the rate theory estimate is in reasonable agreement with the experimental data. Bagchi and co-workers report that their rate estimates are quite sensitive to their chosen coordinates,^7^ making it important to be able to evaluate coordinates with respect to their ability to describe the dynamics objectively. A principled way of doing so is to compare coordinates to the committor,^23^ which is the probability that from a given microscopic state a trajectory goes to a product state before a reactant state.^6,24^

Here, we use a generalization of MSMs termed dynamical Galerkin approximation (DGA)^25,26^ to estimate the committor for insulin dimer dissociation. We show that the committor correlates with the number of interfacial contacts (consistent with the choices of CVs in Refs. 15 and 17) and the total number of interfacial water molecules. We use the committor, together with the equilbrium probability, to compute the reactive current, which enables us to quantify the importance of competing pathways and estimate the rate of dissociation. We show that the heterogeneous paths out of the dimer state and the fluxes between them can be understood in terms of three axes around which the monomers rotate, providing a simple perspective on this complex reaction.

## Methods

Here we describe the statistics that we use in our analysis and how we compute them. In brief, we run many short, unbiased simulations and use them to define a set of Markov states and the probabilities of transition between them. Because we draw the initial structures for these simulations from previous equilibrium simulations, we can use the earlier results to estimate the equilibrium probability (*π*) over the states, which provides information about relative stabilities of the Markov states. From the matrix of transition probabilities, we can compute the committor (*q*), which, as defined above, is the probability of reaching a product state before a reactant state from a given Markov state. Because it provides information about the likelihood of completing a stochastic reaction, the committor serves as a measure of reaction progress, and we define the transition state as the ensemble of states with equal likelihood of next going to the reactant (*A*) and product (*B*) states (*q*_*A*_ = *q*_*B*_ = 0.5). From the equilibrium probability and the committor, we can compute the reactive current, which tracks how probability flows within the ensemble of reactive trajectories; in turn, we integrate over the reactive current along the committor to obtain the reactive flux (*R*_*AB*_; we denote the fractional contribution to the flux of state *i* by *F*_*i*_). The reactive flux divided by the fraction of time the system spends having last come from state *A* is the rate constant (*k*_*AB*_). Figure 2 summarizes our computational workflow.

**Figure 2.**
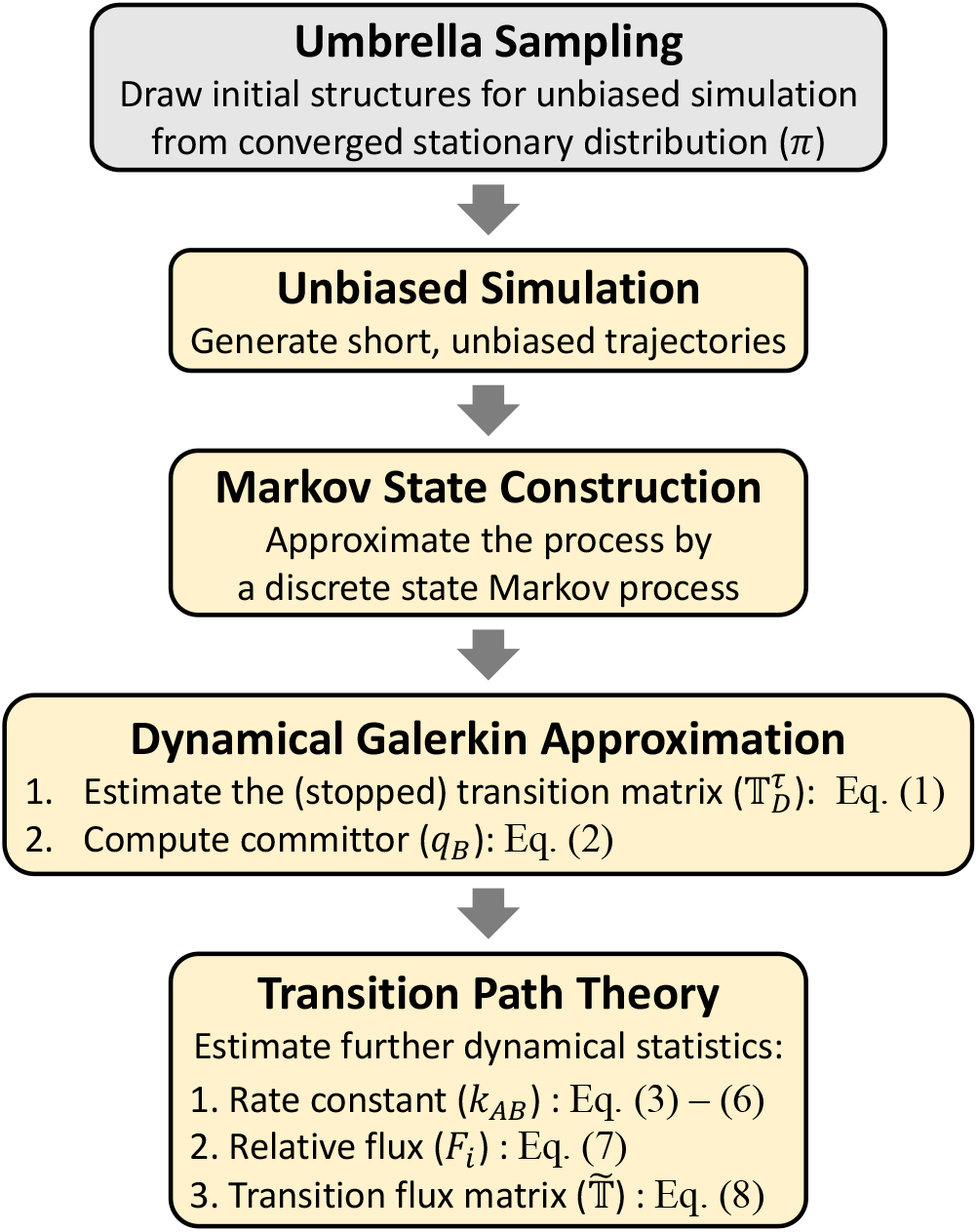
Schematic of the workflow. Gray box shows calculations performed in Ref. 15, and yellow boxes show calculations in the present study.

### Simulation data

In principle, both *π* and *q* can be obtained from DGA using an arbitrary sample distribution.^25–28^ We initially generated a sample distribution by running unbiased dynamics from conformations sampled by adiabatic-bias molecular dynamics (ABMD),^29^ following previous studies.^26,30^ However, we found that the resulting PMF was inconsistent with that obtained from earlier replica-exchange umbrella sampling simulations (REUS),^15^ which were converged with the aid of an error estimator.^31^ Therefore, we instead selected conformations from the umbrella sampling simulations as the starting points for unbiased dynamics simulations, as we describe below.

The system setup was the same as in the REUS simulations, and we refer the reader to Ref. 15 for details. To summarize, the system was constructed from a crystal structure of human insulin (PDB ID 3W7Y^13^) and modeled in aqueous solution with cubic periodic boundary conditions. The protein, 48 K^+^ ions, and 44 Cl^−^ ions were represented by the CHARMM36m^32^ force field, and 15,532 water molecules were represented by the TIP3P model;^33^ the total number of atoms was 48,260. After equilibration, the simulation box size was fixed at (7.82 nm)^3^.

From each of the 748 REUS simulations, we sampled 24 structures equally distributed along the 5 ns trajectory, resulting in 24 *×* 784 = 18,816 initial structures. From each sampled structure, we initialized a 5 ns trajectory using OpenMM 7.7,^34^ yielding an aggregate simulation time of 94.08 μs. Simulations were performed in the canonical (isothermal-isochoric) ensemble at 303.15 K using the LFMiddle integrator^35^ with a time step of 2 fs and a friction constant of 0.083 ps^−1^. The particle-mesh Ewald method was used to calculate non-bonded forces with a cutoff distance of 1.2 nm; the non-bonded interactions were smoothly switched off from 1.0 to 1.2 nm through the built-in OpenMM force-switch function. The lengths of bonds to hydrogens were constrained using the SHAKE algorithm.^36^ Structures were saved every 5 ps.

### Markov state model (MSM)

We used the simulation data described above to define discrete states and estimate the probabilities of transition between them. In this section, we describe variables that we use to characterize the system and states that we define in terms of them.

#### Definition of the dimer and monomer states

We used the following variables to define the dimer and (separated) monomer states based on previous studies:

1. RMSD_int_ measures the root mean square deviation (RMSD) of interfacial C^*α*^ atoms from their positions in the crystal structure. We define the interfacial C^*α*^ atoms to be those within 10 Å of any C^*α*^ atom of the other monomer in the crystal structure.^18^
2. Φ_*α*_ is a pseudodihedral angle that measures twist of the interfacial *α*-helices.^15,20^ It is defined by the geometric centers of Ser^B9^–Leu^B11^, Val^B12^–Tyr^B16^, Val^B′12^– Tyr^B′16^, and Ser^B′9^–Leu^B′11^ (the primes distinguish one monomer from the other).
3. 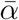 measures the separation of the interfacial *α*-helices.^15^ It is the average distance between C^*α*^ atoms of the following pairs of residues: Ser^B9^-Tyr^B′16^, Ser^B9^-Glu^B′13^,Val^B12^-Tyr^B′16^, Glu^B13^-Glu^B′13^, Glu^B13^-Ser^B′9^, Tyr^B16^-Val^B′12^, and Tyr^B16^-Ser^B′9^.
4. 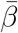 measures the separation of the interfacial *β*-strands.^15^ It is the average distance between C^*α*^ atoms of the following pairs of residues: Phe^B24^-Tyr^B′26^, Phe^B25^-Phe^B′25^, and Tyr^B26^-Phe^B′24^.
5. *N*_c_ measures the number of pairs of non-hydrogen atoms from separate monomers that are within 4.5 Å of each other.
6. *N*_sw_ measures the number of shared waters. The shared waters are ones with oxygen atoms that are simultaneously within 4 Å of at least one interfacial non-hydrogen atom from each monomer. Here, we define the interfacial atoms to be those within 4 Å of a non-hydrogen atom from the other monomer in the crystal structure.

Given these features, we define the dimer state (*A*) as structures with RMSD_int_ *<* 2 Å, 120^*°*^ *<* Φ_*α*_ *<* 135^*°*^, 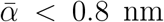, and 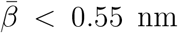. We define the monomer state (*B*) as structures with RMSD_int_ *>* 10 Å, *N*_c_ = 0, and *N*_sw_ = 0.

#### Features used to define states

We divide the remainder of the conformation space into *k* − 2 states so that there are *k* states with the dimer and monomer states. Unless otherwise indicated, we use *k* = 600. We discuss the choice of the number of states in the Supporting Information (text and Figure S2). We cluster the data with *k*-means using 45 features: 26 local features and 19 global features.

The local features correspond to contacts between 26 residue pairs (Table S1 and Figure S1). These are based on the residues used to define 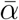 and 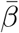, as well as residues in the *β*-turn and close to the C-terminus. We consider more residues in B21–B29 than in the *α*-helices because this segment is more heterogeneous as dissociation proceeds. For each residue pair, we compute the distance between the C^*α*^ atoms, *x*_*i*_, and then take as the feature tanh((*x*_*i*_ − *μ*_*i*_)*/*2*σ*_*i*_), where *μ*_*i*_ and *σ*_*i*_ are the mean and standard deviation of *x*_*i*_ over the entire dataset.

We found that the local features were insufficient to characterize conformations at later stages of dissociation, when most contacts are broken. Consequently, we also include global features. The global features consist of half the distance between the monomer centers of mass, *R*_com_*/*2, and the sine and cosine of nine angles:

1. Two pseudodihedral angles, Φ_*α*_ and Φ_*β*_, measuring twist of the interfacial *α*-helices and *β*-strands, respectively; Φ_*α*_ is defined above, and Φ_*β*_ is defined by the C^*α*^ atoms of Tyr^B26^, Phe^B25^, Phe^B^*′*^25^, and Tyr^B^*′*^26^;
2. Two angles, *γ* and *γ*^*′*^, which measure splaying of the interfacial *α*-helices (see Figure 7 below); *γ* is the angle defined by the geometric centers of Val^B12^–Tyr^B16^, Phe^B25^, and Phe^B^*′*^25^, and *γ*^*′*^ is the corresponding angle for the other monomer;
3. Five Euler angles characterizing the relative orientations and rotations of the monomers.^37^

As defined above (Table S1), the different types of features have comparable magnitudes, and we use them in the *k*-means algorithm without further scaling. Because trajectories tend toward the stable states, we used only the 50 frames from 255 ps–500 ps of the 5 ns trajectories when defining the cluster centers to ensure sufficient resolution outside the stable states. We used MDAnalysis^38,39^ to compute all collective variables.

### Estimation of equilibrium statistics

As a consequence of the fact that we initialize the unbiased simulations from previous umbrella sampling simulations, the equilibrium probabilities of the conformations sampled by the unbiased simulations are proportional to the weights, *w*(**x**), of their associated initial conformations. After normalizing these weights so they sum to one, we compute the PMF over Markov states as

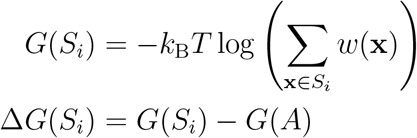

where *S*_*i*_ denotes Markov state *i*, **x** is a structure in *S*_*i*_, *k*_B_ is Boltzmann’s constant, and *T* is temperature.

### Estimation of dynamical statistics

From the MSM, we compute the committor, reactive current, and rate. The central quantity in an MSM is the matrix of state-to-state transition probabilities (𝕋). A key aspect of DGA that distinguishes it from a traditional MSM is that the transition transition matrix is constructed with statistic-specific boundary conditions. That is,

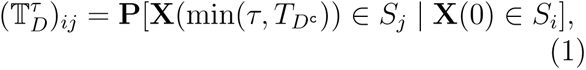

where **X**(*t*) is the structure at time *t, S*_*i*_ and *S*_*j*_ are Markov states, 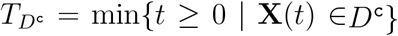 is the time of first exit from the domain *D* (here, *D* = (*A ∪ B*)^c^, but we consider other choices below), and *τ* is a lag time that must be chosen to be sufficiently long that the dynamics are approximately Markovian in the variables used to define the states. The minimum in (1) effectively stops trajectories when they enter states *A* and *B*. We do not symmetrize 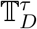to enforce microscopic reversibility.

We denote the committor that is the probability of next going to state *B* rather than state *A* from structure **x** by *q*_*B*_(**x**). By definition, *q*_*A*_ = 1 − *q*_*B*_, and by microscopic reversibility, *q*_*A*_ is also the probability of last coming from state *A* rather than state *B*. The committor satisfies the equation

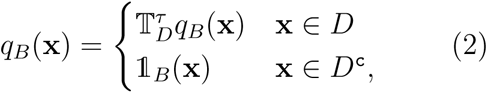

where 𝟙_*B*_(**x**) is an indicator function that is one if **x** is in state *B* and zero otherwise. We solve for *q*_*B*_ with a modified algorithm that treats non-Markovian effects through the addition of memory terms.^40^ Here, we show committors estimated using a lag time of 2 ns and 1 memory term, which we choose to balance expressivity and statistical error (see discussion in Supporting Information and Figures 4, S2, and S3).

Given *π, q*_*A*_, and *q*_*B*_, we use transition path theory (TPT)^41^ to compute the reactive current projected onto a vector of CVs, *θ*:

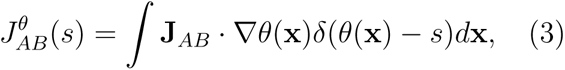

where

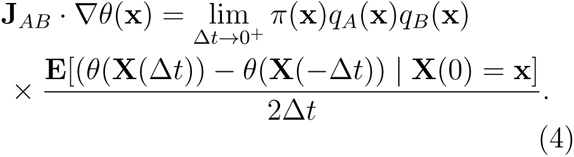

Integrating the reactive current along the committor (i.e., setting *θ* = *q*_*B*_ in (3) and (4)) gives an estimate of the number of reactive trajectories per unit time, *R*_*AB*_, or the net reactive flux:

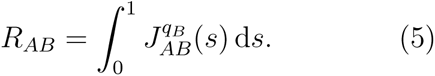

Dividing *R*_*AB*_ by the fraction of time the system spends having last come from state *A* gives the TPT rate constant:

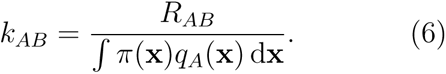

In practice, we use finite-lag time estimators for 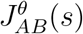, *R*_*AB*_, and *k*_*AB*_ ^26,42^—we translate these estimators to our notation and provide further details concerning their use in the Supporting Information.

To estimate the contribution of a state to the net reactive flux, we compute the dynamical statistics with an extra state excluded from the domain,^43^ *D* := (*A ∪ B ∪ C*_*i*_)^c^ where *C*_*i*_ is any (coarse-grained) state of interest. The interpretation of the committor 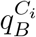 then becomes the probability to next go to state *B* rather than *A ∪ C*_*i*_, and 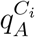 becomes the probability to last come from state *A* rather than *B ∪ C*_*i*_. We can solve for the committor 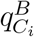 using (2) with *D* := (*A ∪ B ∪ C*_*i*_)^c^. Similarly, we can solve for 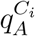 using the same equation but with *A* and *B* switched. We then compute the number of reactive trajectories from *A* to *B* without entering *C*_*i*_ per unit time, which we label 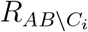, by inserting 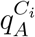 and 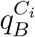 into (4) and (5). We then define the fractional contribution of each coarse-grained state to the net flux as

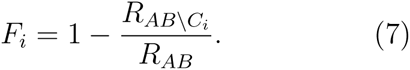

The procedure above can be viewed as a simplified version of the approach for computing history-dependent TPT statistics in Ref. 44.

In the **Results**, we aggregate Markov states into eight coarse-grained states. To systematically evaluate how the coarse-grained states evolve, we compute a coarse-grained transition flux matrix, 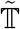, where 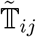 is the probability for a structure starting in state *C*_*i*_ to go to *C*_*j*_ without visiting any other states. More specifically, the elements of 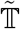 are

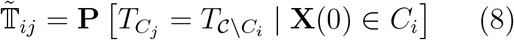

where

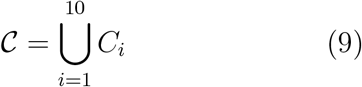

and *C*_*i*_ represents the dimer, monomer, and coarse-grained states.

## Results

Our analysis of insulin dimer dissociation is organized as follows. We first present the PMFs, committor, and reactive currents as functions of the CVs used to control the sampling. While we project these quantities to two dimensions for visualization, we compute them for 600 Markov states defined in 45 dimensions, as described in **Methods**, and we use them without projection to estimate the rate of dissociation. To dissect the mechanism, we identify CVs that correlate with the committor and relate them to the dimer structure. Then, we consider CVs that do not correlate with the committor to characterize the diversity of reaction paths and their associated fluxes.

### The inverse rate is approximately 100 ms

As described in **Methods**, we draw the initial conformations for our unbiased molecular dynamics simulations from umbrella sampling simulations with restraints in two CVs: the average occupancies of seven contacts between the interfacial *α*-helices 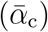 and three contacts between the interfacial *β*-strands 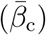.^15^ We show the PMF, committor, and reactive currents as functions of these CVs in Figure S4 and on the average distances for the atom pairs defining these CVs (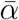 and 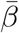) in Figure 3.

**Figure 3.**
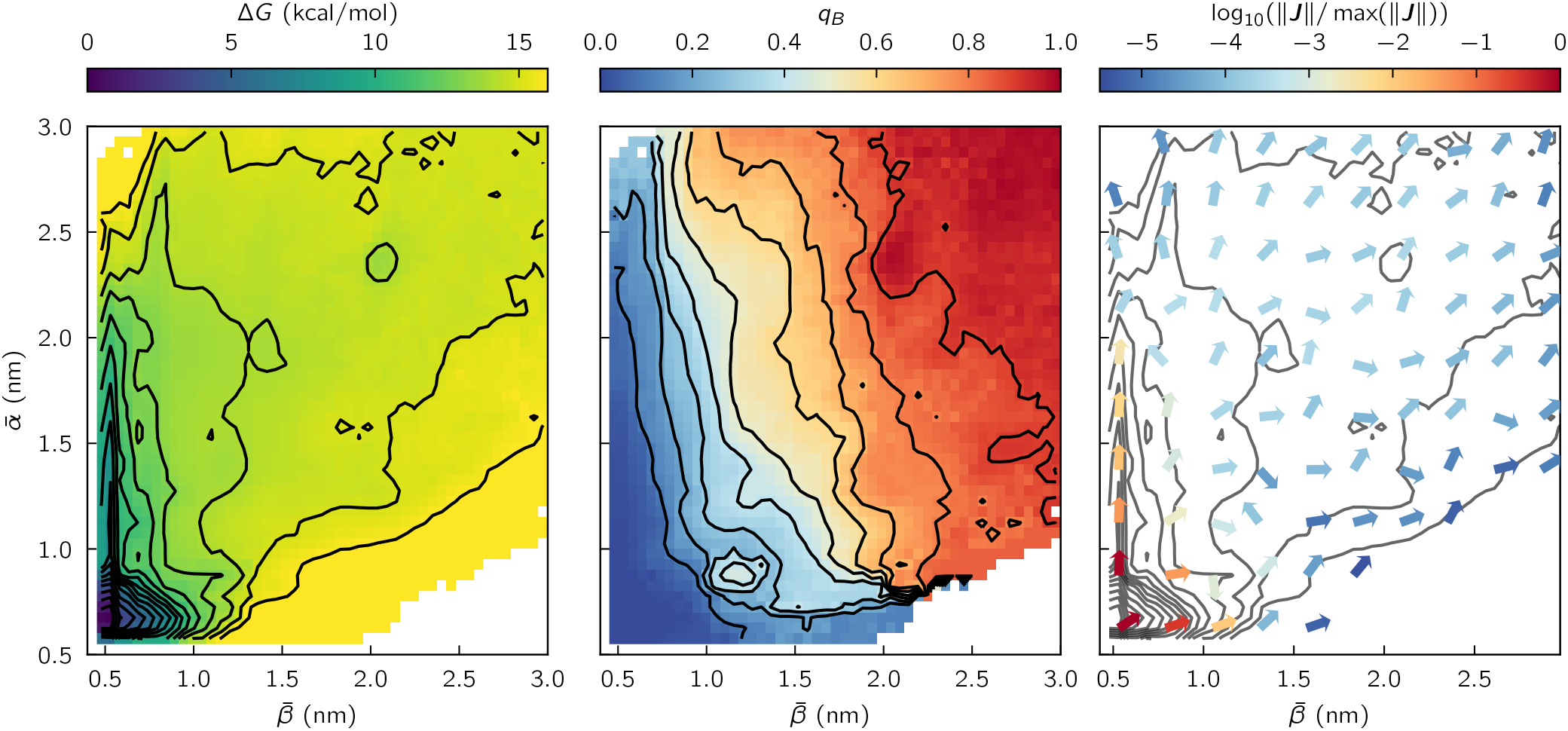
Projections of reactive statistics. The potential of mean force Δ*G* (left), committor *q*_*B*_ (middle), and reactive current 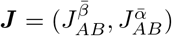 (right, arrows; contours reproduce Δ*G* from left) as functions of 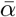 and 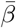. (left and right) Contours are drawn every 1 kcal/mol.(middle) Contours are drawn every 10% increase of probability.

**Figure 4.**
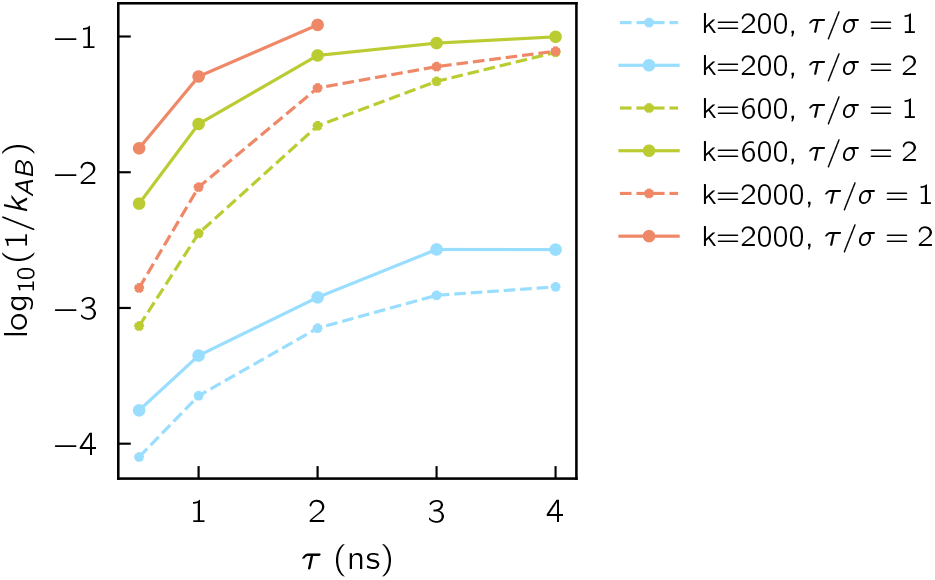
Dependence of inverse reaction rate estimates (in s) on hyperparameters: the number of Markov states, *k*, the lag time, *τ*, and the number of memory terms, *τ/σ* − 1, where *σ* is the time interval used to compute the memory terms.

The DGA/MSM PMFs closely resemble those from umbrella sampling,^15^ indicating that the initial conformations are indeed consistent with the equilibrium distribution. Focusing on Figure 3, the committor increases smoothly from the dimer state in the lower left corner to the monomer state in the upper right corner. The lines of constant committor values (isocommittors) are diagonal, and the arrows representing the reactive currents fan out from the dimer state. Both of these features support the idea that the reaction can follow diverse paths, with the interfacial *α*-helices or *β*-strands separating in either order or concomitantly.

We integrate the flux along the committor to estimate the rate, as described in Methods. As we vary the lag time, the estimate appears to converge to the order of 10 s^−1^ for the rate, or 100 ms for its inverse (Figure 4), markedly slower than both experimental temperature-jump studies performed in ethanol^21,22^ and previous computational estimates.^7^ We discuss these studies further in the **Discussion**.

As a first step toward understanding the factors that contribute to the rate, we plot the potential of mean force as a function of the committor in Figure 6a (top left). There is a barrier of about 13 kcal/mol at *q*_*B*_ *≈* 0.14 (blue shading), and the potential of mean force is relatively flat between *q*_*B*_ *≈* 0.14 and *q*_*B*_ *≈* 0.9. We relate these features to CVs in the following section, where we also describe how information is represented within the plots in Figure 6a.

### The committor correlates with the total number of interfacial contacts

With a view toward structurally interpreting the committor, we computed its correlation with a large number of CVs (Figure 5).^23,27^ We found that the CVs used to control sampling in previous studies 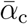 and 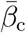 in Ref. 15 and the center-of-mass distance, *R*_com_, and number of C^*α*^ intermonomer contacts, *Q*, in Ref. 17 are highly correlated with the committor. In addition to those CVs (and the closely related CVs 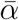 and 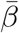), the CVs with the highest correlation include the total number of intermonomer contacts, *N*_c_, the total number of water molecules at the dimer interface, *N*_w_, which increases as the dimer interface becomes exposed, and the number of shared water molecules at the dimer interface, *N*_sw_, which decreases as the monomers separate.

**Figure 5.**
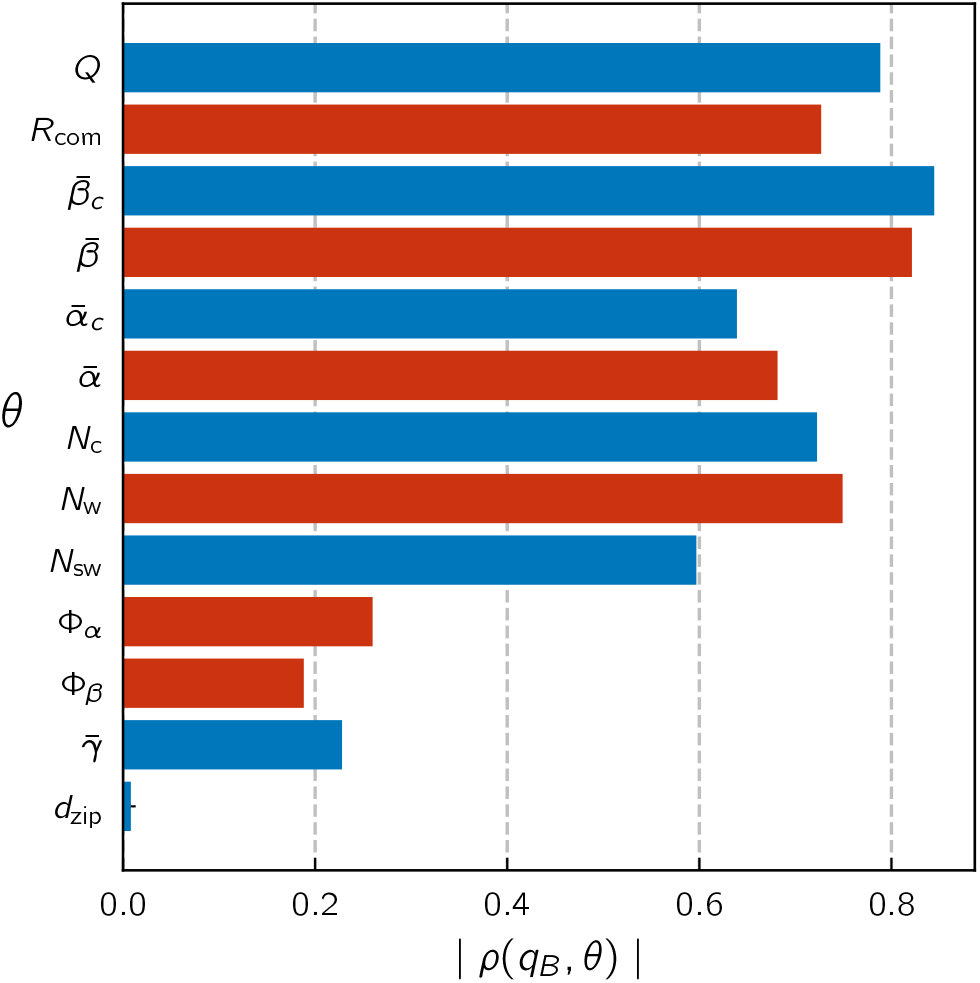
Pearson correlation coefficient between the committor and physical CVs of interest over the entire data set (18,816,000 structures). Red and blue colors correspond to positive and negative correlation coefficients, respectively.

**Figure 6.**
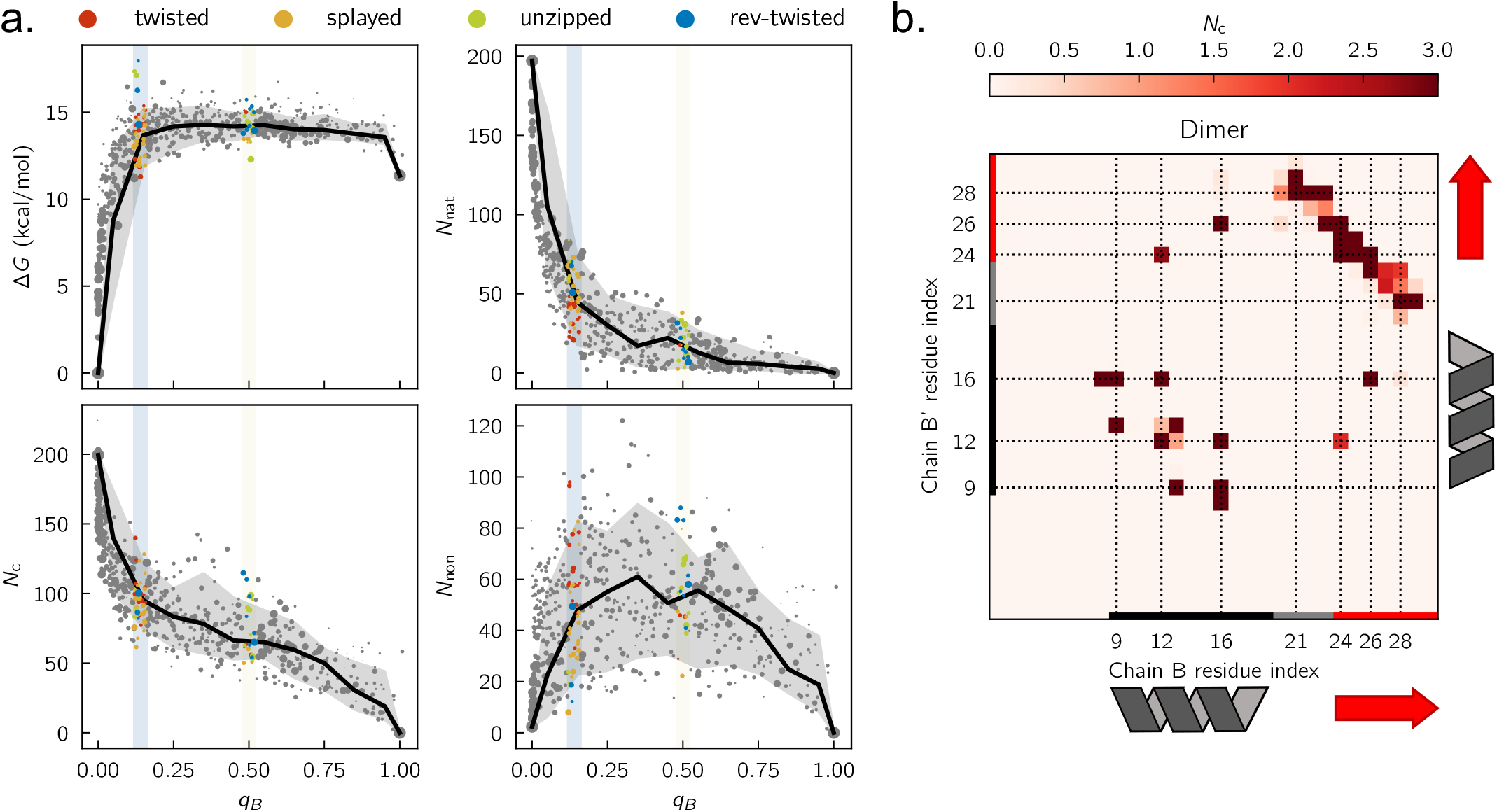
Comparison of the free energy profile to the evolution of the number of contacts. **a.** CVs as functions of the committor: (top left) free energy, (bottom left) total number of intermonomer contacts, (top right) native contacts, and (bottom right) nonnative contacts. The size of each symbol is proportional to the reactive flux passing through the associated Markov state. Blue and yellow bars mark *q*_*B*_ *≈* 0.14 and *q*_*B*_ *≈* 0.5. Symbol colors at *q*_*B*_ *≈* 0.14 and *q*_*B*_ *≈* 0.5 indicate coarse-grained state assignments. We divide the horizontal axes into intervals of 0.1 and sort the states within each interval by their vertical axis values; the black line marks the value corresponding to 50% of the flux, and the gray shaded region shows the [10%, 90%] interval of the flux (i.e., 40% on either side of the line). **b**. Average residue-residue contact map of the dimer state. See Figure S7 for the average atom-atom contact map.

**Figure 7.**
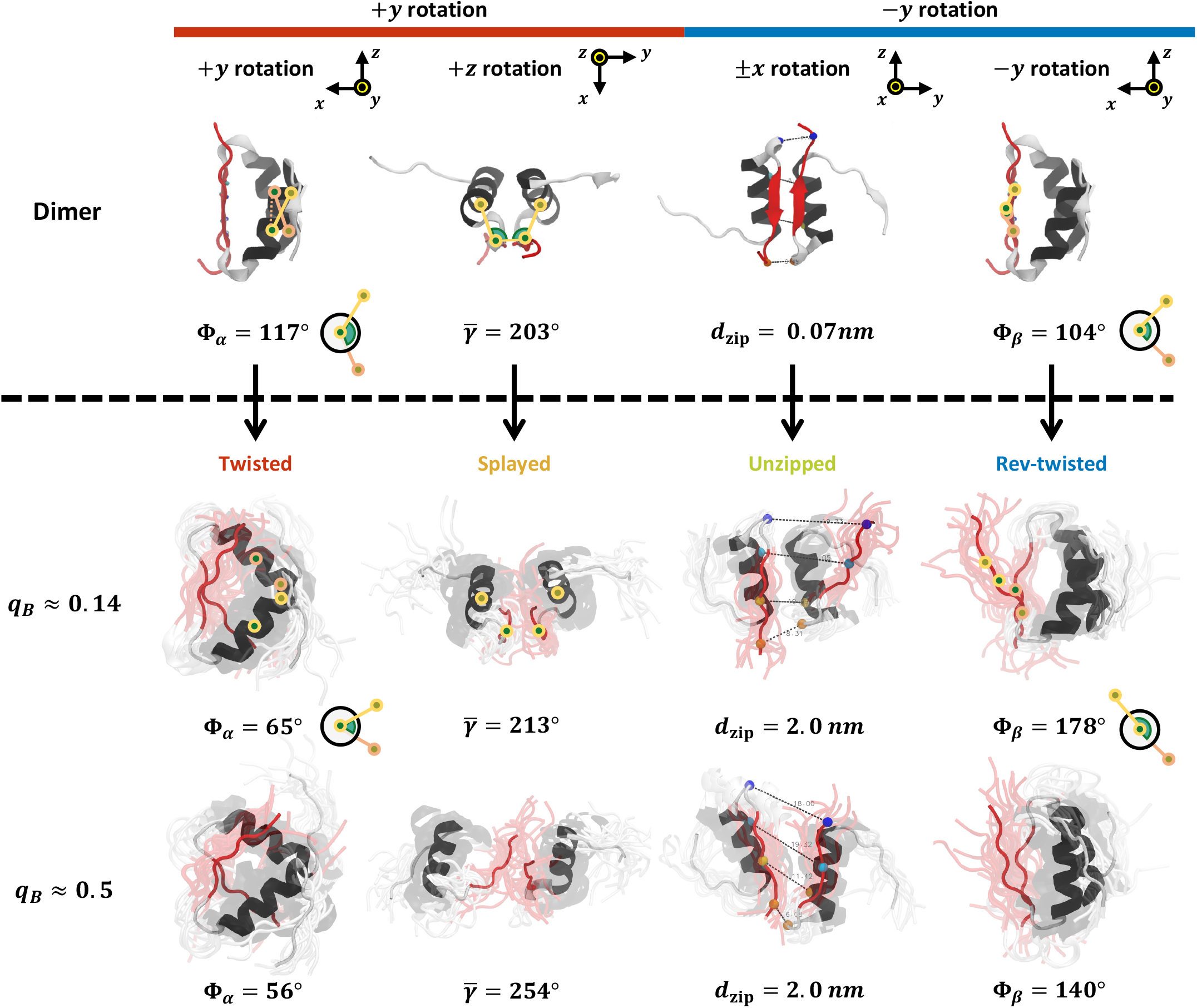
Coarse-grained states. (top row) Collective variables used to define the states and their associated rotations. (bottom two rows) Representative structures at *q*_*B*_ *≈* 0.14 and *q*_*B*_ *≈* 0.5 as indicated. To obtain the structures shown, we randomly sampled ten structures (translucent) from each of the Markov states associated with each coarse-grained state and then iteratively aligned the structures to their average structure until the backbone RMSDs to the average structure (opaque) converged. See Figures S9 and S10 for alternative views of the coarsegrained states.

Although the correlation coefficient is useful for identifying CVs that track monotonically with the committor, it can obscure nonmonotonic relationships, as well as more subtle relationships (e.g., how quickly a variable changes as the reaction progresses). Consequently, we also show plots in which we plot the average value of a variable for each Markov state as a function of its committor value; we term these evolution profiles. To highlight Markov states with the highest fluxes, we make the symbol sizes proportional to the fluxes, and we also indicate the flux-weighted quantile of the variable for the Markov states at each value of the committor (using bins of 0.1; see Figures 6 and S5). Figure 6a shows the evolution profile for *N*_c_, which is strongly correlated with the committor. There are two phases: there is a relatively rapid drop in *N*_c_ from the dimer state to *q*_*B*_ *≈* 0.14, followed by a slow, steady decrease to zero at *q*_*B*_ = 1. The trends in the *N*_c_ profile are mirrored in the *N*_w_ profile (Figure S5); the two CVs track each other almost perfectly because water molecules fill the space vacated by protein (Figure S6). The initial rapid phase in the *N*_c_ profile coincides with the rise in free energy, while the subsequent slow phase coincides with the plateau in free energy.

To dissect this behavior, we separate the intermonomer contacts into native and nonnative ones. To determine the native contacts, we examine the percentage of structures in the dimer state that each (atom-atom) contact is occupied. Because we find there is no natural cutoff for these occupations, we take the native contacts to be those occupied in at least 1% of the structures in the dimer state (Figures 6b and S7; other cutoffs gave qualitatively similar results). We see that the drop in *N*_c_ from the dimer state to *q*_*B*_ *≈* 0.14 results from an even more rapid drop in native contacts offset by a rise in nonnative contacts. While most of the rise in nonnative contacts is in this initial phase, the number of nonnative contacts peaks around *q*_*B*_ *≈* 0.5, which we define as the transition state since states with this committor value have an equal probability of next going to the dimer and monomer states. Thus, the transition state is an encounter complex with about 20 native contacts and 50 nonnative contacts. However, this simple description masks structural diversity that we elaborate in the next section.

### The diversity of pathways can be understood in terms of three axes of rotation based on the dimer structure

Given that the CVs that track closely with the committor are compatible with a diversity of structures at each stage of the reaction^15^ (Figures 3, S4, and S5), we wanted to determine whether insulin dissociates by distinguishable pathways, or the reactive path ensemble is better viewed as a continuum. Answering this question and, more generally, developing a structural understanding of the mechanism can aid in interpreting the effects of mutations and designing new insulin analogues, as well as connecting our results to structural ideas about molecular recognition.^15,30,46^ To this end, we sought to identify structural CVs that are orthogonal to the committor and to use them to characterize the reactive path ensemble.

Based on the results of previous studies,^15,17,20^ we considered CVs that track twisting of the interfacial secondary structure elements. The twisting of the *α*-helices relative to each other can be quantified by a pseudodihedral angle, Φ_*α*_, defined by the geometric centers of the backbone atoms of segments Ser^B9^– Leu^B11^, Val^B12^–Tyr^B16^, Val^B′12^–Tyr^B′16^, and Ser^B ′9^–Leu^B′11^. Similarly, the twisting of the *β*-strands relative to each other can be quantified by a pseudodihedral angle, Φ_*β*_, defined by the C^*α*^ atoms of Tyr^B26^, Phe^B25^, Phe^B′25^, and Tyr^B′26^. Note that these definitions differ slightly from those in Ref. 15, so that the dimer state corresponds to the *trans* state of the pseudodihedral angle in each case. These CVs are correlated (*ρ*(Φ_*α*_, Φ_*β*_) = 0.65; Figure S8) and characterize rotations around axes that are orthogonal to the dimer interface, parallel to the line connecting the centers-of-mass of the monomers (labeled the *y*-axis in Figure 7).

Motivated by this observation, we defined two other axes based on the dimer structure. We defined the *x*-axis as the *C*_2_ rotation axis of the dimer and the *z*-axis as the cross product of the *x*- and *y*-axes, which is parallel to the *β*-strands. We then defined CVs that characterize rotations around the *x*- and *z*-axes. Because the monomers are not rigid bodies, one might expect more than one variable to be required to define the dynamics around each axis (as is the case for the *y*-axis, where we track both Φ_*α*_ and Φ_*β*_). Nevertheless, we found that only one additional CV for each axis was sufficient. To describe the rotation around the *z*- axis, we define three angles: *γ* is the angle of the geometric centers of the segments Val^B12^– Tyr^B16^, Phe^B25^, and Phe^B^*′*^25^; *γ*^*′*^ is the corresponding angle for the other monomer; and 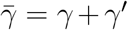. To describe the rotation around the *x*-axis, we combine four C^*α*^-C^*α*^ distances: *d*_zip_ = (*d*_B21−B_*′*_29_ + *d*_B16−B_*′*_26_) − (*d*_B29−B_*′*_21_ + *d*_B26−B_*′*_16_). Figure 7 illustrates the axes and the CVs that characterize rotation around them.

We clustered the Markov states into coarse-grained states based on Φ_*α*_, Φ_*β*_, 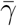, and |*d*_zip_|. By varying the number of clusters, we found that four coarse-grained states were sufficient to capture the qualitative features of the pathways. Each is distinguished from the others by a shift in one of the CVs relative to its value in the dimer state: the twisted state has low Φ_*α*_, the splayed state has high 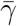, the unzipped state has high |*d*_zip_|, and the rev-twisted state has high Φ_*β*_ (Figure S8). However, we emphasize that each such shift can be accompanied by others owing to correlations between the CVs.

Figures 7, S9, and S10 show structures of the coarse-grained states at different stages of the reaction, Figures 8 and S7 show their contact maps, and Figure S11 shows their solvation. At *q*_*B*_ *≈* 0.14, both the twisted and splayed states exhibit a rotation in the positive direction around the *y*-axis and in turn relatively low Φ_*α*_ and Φ_*β*_ (Figure S8; see Figure S9 for an alternative view of the splayed state that makes clear the +*y*-rotation). This rotation shifts contacts of the C-terminal residues (B26– B30) of each monomer from the *β*-turn (B^*′*^21) to the C-terminal end of the *α*-helix (B^*′*^16) of the other monomer in a symmetric fashion. By contrast, the unzipped and rev-twisted states show detachment of the B^*′*^26–B^*′*^30 residues of one monomer (in Figures 7 and 8 we show data only for states with *d*_zip_ *>* 0 to make this asymmetry apparent; states with *d*_zip_ *<* 0 are similar but the monomers are switched).

**Figure 8.**
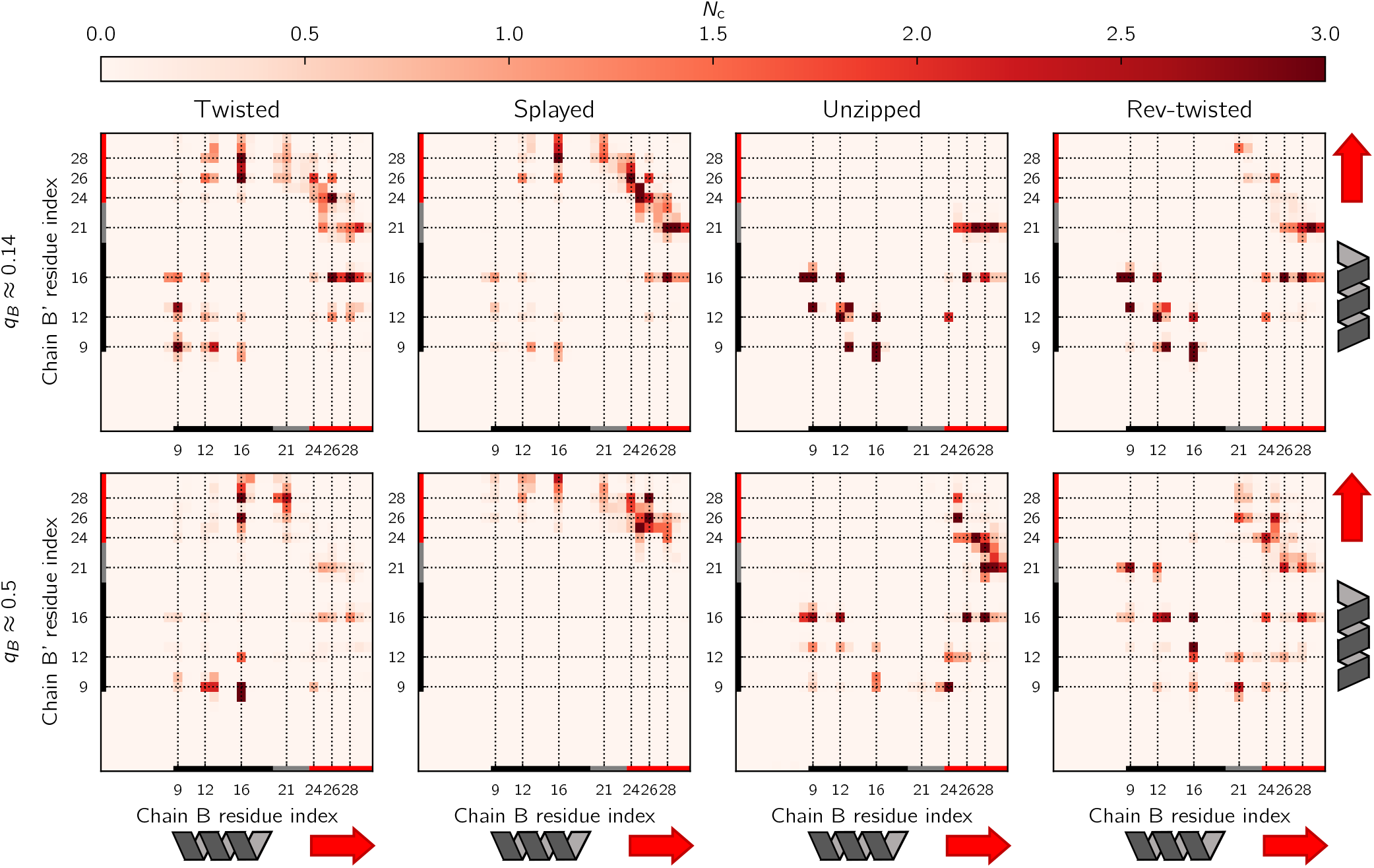
Average residue-residue contact map of each of the coarse-grained states. See Figure S7 for the average atom-atom contact map.

The twisted and splayed states are differentiated by whether they exhibit further rotation around the *y*-axis or rotation around the *z*-axis. In the twisted state, further rotation around the *y*-axis leads to contacts between B26–B30 and B^*′*^12–B^*′*^13 in the interfacial *α*- helix of the other monomer, and it brings the N-termini of the interfacial *α*-helices together to make a B9–B^*′*^9 contact. In the splayed state, rotation around the *z*-axis results in a loss of contacts between the interfacial *α*-helices. Similarly, the unzipped state and rev-twisted states (which again occupy the same region of the Φ_*α*_Φ_*β*_-plane; Figure S8) are differentiated by the rotation around the *x*-axis. In the unzipped state, this rotation results in a complete loss of contacts for the detached B^*′*^26–B^*′*^30 residues at *q*_*B*_ *≈* 0.14.

At *q*_*B*_ *≈* 0.5, the coarse-grained states are more heterogeneous. In the twisted state, only contacts between the *β*-turn region (B16 and B21) and the C-terminus (B^*′*^26–B^*′*^30) or the N-terminus of the *α*-helix (B^*′*^9) of the other monomer persist. In the splayed state, the contacts between the interfacial *α*-helices are completely lost, and one of the two groups of contacts between the C-terminal residues (B26– B30) and the *β*-turn (B21) is broken while the other remains intact, mirroring the effects of the *x*-rotation discussed above for the unzipped state at *q*_*B*_ *≈* 0.14. Notably, both the twisted and splayed states show asymmetric contact maps.

In the unzipped state at *q*_*B*_ *≈* 0.5, the C-terminus (B26–B30) of one monomer (left in Figure 7) inserts between the *β*-trun side of *α*-helix (B^*′*^16) and that of *β*-strand residues (B^*′*^24) of the other monomer, bringing B28–B30 in contact with the *β*-turn (B^*′*^21–B^*′*^23) as well.

The other monomer’s C-terminus (B^*′*^24–B^*′*^28) is heterogeneous in conformation but samples contacts with B25 of the former. The rev-twisted state rotates the opposite way around the *y*-axis as the twisted state, and this results in a nonnative contact between the C-termini of the interfacial *α*-helices at B16–B^*′*^16 (in contrast to the B9–B^*′*^9 contact for the twisted state at *q*_*B*_ *≈* 0.14) as well as nonnative contacts that arise from the *β*-turn (B21) of one monomer inserting between the C-terminal part of the *β*-strand (B^*′*^26-B^*′*^28) and the N-terminal part of the *α*-helix (B^*′*^9 and B^*′*^12) of the other monomer. Notably, the contact map of the rev-twisted state at *q*_*B*_ *≈* 0.5 is symmetric.

### There is extensive mixing between selected pathways

To quantify the contributions of the pathways, we computed the fractional fluxes through the coarse-grained states as described in Methods (Figure 9a). At *q*_*B*_ *≈* 0.14, the coarse-grained states that involve +*y* rotation (splayed and twisted) make larger contributions to the net flux than the coarse-grained states that involve −*y* rotation (unzipped and rev-twisted). This reflects the fact that the former’s free energies are lower (compare the red and orange symbols with the blue and green ones at *q*_*B*_ *≈* 0.14 in Figure 6a). At *q*_*B*_ *≈* 0.5, the ordering is reversed, presumably owing to mixing between pathways.

**Figure 9.**
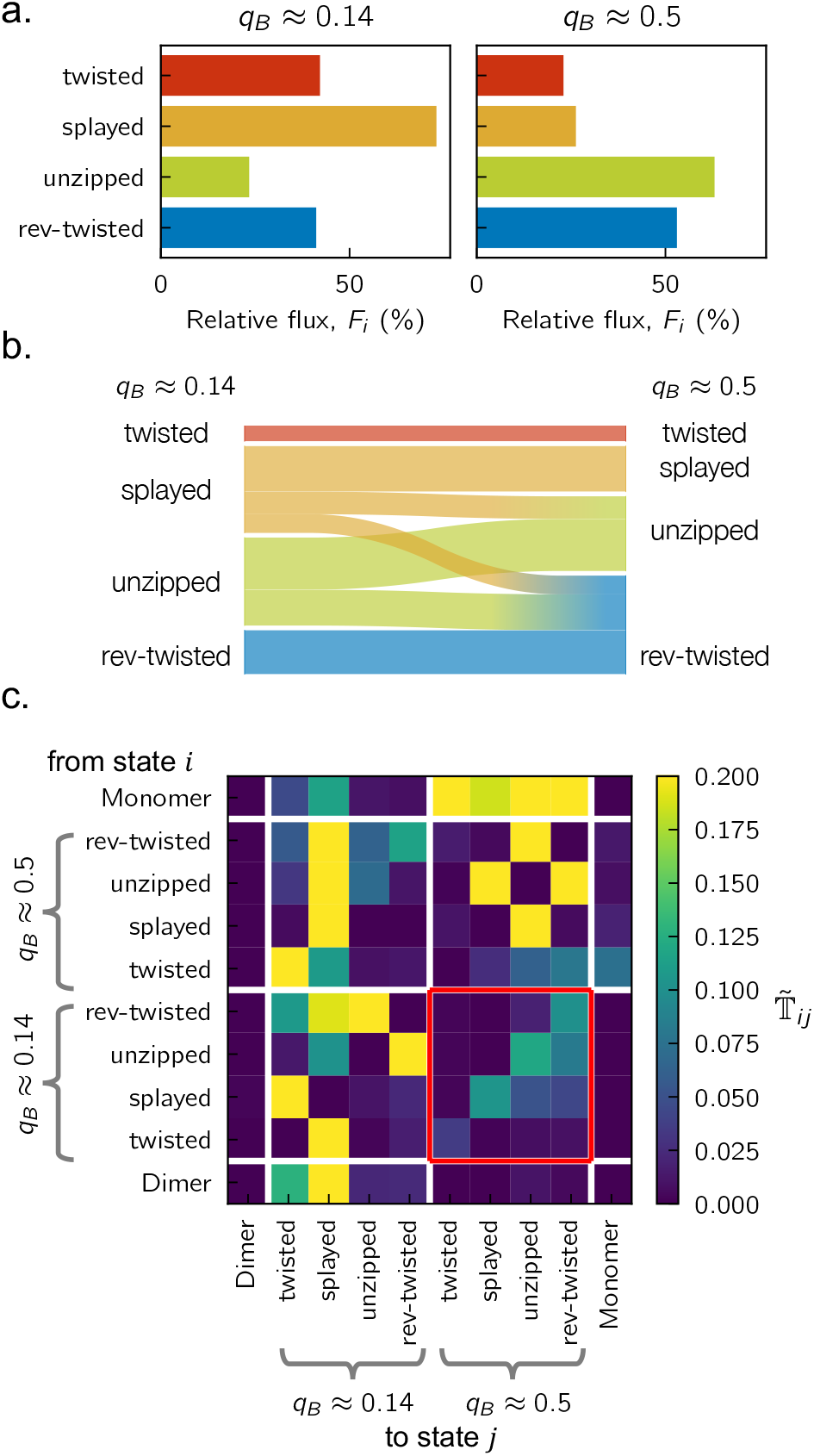
Quantifying the contributions of pathways and their mixing. **a.** Fractional contribution of the coarse-grained states to the net flux. **b**. Sankey diagram of mixing between coarse-grained states. The width of each bar is proportional to the corresponding element in the transition flux matrix 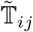. **c**. Transition flux matrix between the coarse-grained states. Red box indicates the entries used to plot the Sankey diagram in **b**.

To characterize the mixing, we computed the coarse-grained transition flux matrix as described in Methods (Figure 9c). One can consider both mixing within each committor value (*q*_*B*_ *≈* 0.14 and *q*_*B*_ *≈* 0.5) and from each to the other. We focus first on the transitions from *q*_*B*_ *≈* 0.14 to *q*_*B*_ *≈* 0.5 (Figure 9b and red box in Figure 9c). The coarse-grained states that involve only rotations around the *y*-axis (twisted and rev-twisted states) flow from *q*_*B*_ *≈* 0.14 to *q*_*B*_ *≈* 0.5 without branching. By contrast, the states that involve multiple rotation axes (splayed state with +*y* and +*z* rotation and un-zipped state with −*y* and +*x* rotation) branch to other states. Furthermore, there is flow from the +*y*-rotation states to the −*y*-rotation states but not vice versa, consistent with the reordering in Figure 9a.

Within *q*_*B*_ *≈* 0.14 (lower left block of Figure 9), states with the same direction of *y* rotation exchange more readily with each other than with states with the opposite direction of *y* rotation. Interestingly, the twisted and splayed states with +*y* rotation at *q*_*B*_ *≈* 0.14 are more likely to proceed to *q*_*B*_ *≈* 0.5 (red box) than to exchange with the unzipped and rev-twisted states with −*y* rotation at *q*_*B*_ *≈* 0.14. On the other hand, the states with −*y* rotation show comparable flux to *q*_*B*_ *≈* 0.5 and exchange with the states with +*y* rotation at *q*_*B*_ *≈* 0.14. At *q*_*B*_ *≈* 0.5 (upper right block of Figure 9), the twisted state appears relatively isolated, while the remaining three states are connected via the unzipped state, which shares structural features with both the splayed state (+*x* rotation) and the rev-twisted state (−*y* rotation).

Put together, our analysis of the structures and fluxes shows that pathways can be distinguished but there is extensive mixing between them, both within each committor value and from one committor value to another. This mixing can be rationalized in terms of the rotation axes that we define based on the dimer structure.

## Discussion

Here, we construct an MSM to characterize insulin dimer dissociation statistically. The heterogeneity of pathways that we document makes this a challenging reaction to study. One key aspect of our approach was initializing the unbiased dynamics simulations used to construct the MSM from structures drawn from converged umbrella sampling simulations.^15,31^ Using a framework that generalizes MSM to solve operator equations with statistic-specific boundary conditions,^25,26^ we computed the equilibrium probabilities and committor values of Markov states and the fluxes through them. From these we obtained a rate and a comprehensive view of the dissociation mechanism.

The estimated timescale of dissociation (approximately 100 ms) is two orders of magnitude slower than the 250–1000 μs timescale ascribed to monomer separation in T-jump studies, which use both elevated temperature and ethanol to accelerate dissociation,^21,22^ in contrast to the simulations. Accordingly, the dissociation constant was shown to decrease by two orders of magnitude in 20% EtOD,^47^ which would be sufficient to account for the difference in *k*_*AB*_ (under the assumption that *k*_*BA*_ does not change). The 2DIR experiments, moreover, measure relaxation of a spectral feature assigned to interfacial *β*-sheet disruption, which would be considered an intermediate state in the simulations.

Our estimated rate is also markedly slower than estimates from Bagchi and co-workers using TST-based theories. The higher barrier that we obtain (compared with their reported values of 7.6–9.3 kcal/mol^7,17^) likely accounts for much of the difference. At the same time, the corrections to TST that they employ to account for dynamics orthogonal to the reaction coordinate are also unlikely to be able to fully capture the dynamics that we describe. Indeed, as the resolution of the model (number of Markov states) becomes lower in Figure 4, we estimate the rate to be faster. TST-based theories applied to a two-dimensional potential of mean force resolve far fewer details than even our coarsest MSM (200 states) and are correspondingly faster still.

We found that the committor correlates closely with CVs that quantify the total number of intermonomer contacts and the total solvent exposure of the dimer interface. Structures in the transition state ensemble, despite their diversity, generally have a relatively small percentage (*<*20%) of native contacts and a highly hydrated interface, consistent with general trends obtained from long, unbiased simulations^18^ and earlier experimental measurements.^1,48,49^ The loss of native contacts at *q*_*B*_ *≈* 0.14 frees the participating residues to make new contacts, and this accounts for the concommitant gain of nonnative contacts (Figure 6a and 8). The fact that we do not see contacts involving residues that are far from the dimer interface supports the idea that encounter complexes can successfully associate only if they first make contact near the native interface,^4,18,^ but our sampling of states that are very different from the dimer may also be incomplete.

We can quantitatively compare the likelihood of association for different encounter complexes based on their contacts. For example, as their committor values indicate, the rev-twisted state at *q*_*B*_ *≈* 0.5 is more likely to dissociate (or less likely to associate) than the twisted state at *q*_*B*_ *≈* 0.14 even though their numbers of native and nonnative contacts are comparable (Figure 6a). This makes clear that details beyond the numbers of native and nonnative contacts are important.

Consistent with this idea, we identified four coarse-grained states that define dissociation pathways. We showed that these pathways can be understood in terms of three orthogonal axes of rotation based on the dimer structure. Along all pathways, the initial event is a rotation around the *y*-axis (Figure 7), consistent with twisted states previously reported.^15,17^ Subsequently, +*z*-rotations can lead to splaying of the interfacial *α*-helices and *x*-rotations can lead to unzipping of the interfacial *β*-strands. These dynamics are consistent with low-frequency normal modes.^19^

Despite the diversity of pathways that we observe, overall, less disorder in the separated monomers than an earlier study of the insulin monomer^10^ and as suggested by experiments.^21,47^ However, these studies employed mutations and/or low pH to limit self-association. The only significant disorder that we observe over the course of the reaction is the detachment of the C-terminal *β*-strand (as measured by 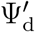, following Ref. 15, and Ψ_B_*′*_25B_*′*_30d_, following Ref. 10; shown in Figures S5 and S12, respectively). In Ref. 15, the detachment angle was averaged over the monomers and appeared to vary more over the *α*-path when projected on 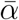 and 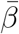. In our case, we resolve the contributions from the monomers. While we see evidence of detachment in both Ψ_d_ and 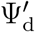 along the pathways involving +*y*-rotation (i.e., the twisted and splayed states; Figure S5), which we associate with the *α*-path, we observe that 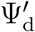(Figure S5) and Ψ_B_*′*_25B_*′*_30d_ (Figure S12) exhibit extreme values at *q*_*B*_ *≈* 0.14 along the pathways involving −*y*-rotation (i.e., the unzipped and rev-twisted states), which we associate with the *β*-path; the latter feature may be obscured when averaged. Interestingly, consistent with Ref. 15, the detachment is more pronounced at intermediate stages of the reaction than in the monomer state.

As the total number of contacts decreases over the course of the reaction, the number of water molecules in contact with the monomers increases. At no point do we see any evidence of a dewetting transition (Figure S6), as observed for some systems,^52,53^ including the insulin dimer.^54^ In the case of the twisted and splayed states, the initial +*y*-rotation brings Lys^B29^ close to the hydrophobic core (Val^B12^, Tyr^B16^, Phe^B24^, and Tyr^B26^, and the corresponding residues from the other monomer); in the case of the unzipped and rev-twisted states, −*y* rotation promotes *β*-strand detachment. Both the presence of charged residues and the absence of an extended hydrophobic surface are expected to suppress dewetting.^55–57^ That said, dewetting may be very sensitive to details of the simulation conditions, and our choice of features may bias our analysis towards solute rather than solvent dynamics;^58^ further analysis of the solvent in our simulations is warranted in the future.

The dynamics of B28-B29 are also important for understanding fast-acting therapeutic insulin analogues. Insulin lispro^59^ (Pro^B28^→Lys and Lys^B29^→Pro) and insulin aspart^60^ (Pro^B28^→Asp) are thought to accelerate dimer dissociation by destabilizing C-terminal native contacts.^61–63^ In our simulations, we observe that these residues make many nonnative contacts as the reaction progresses, suggesting that a full understanding of the effects of these mutations on the kinetics requires consideration of nonnative interactions as well. A further complication is that mutations can shift the importance of competing pathways, as previously shown for dissociation of phenol from the insulin hexamer.^30^ We expect such a shift to be possible in the case of dimer dissociation given the extensive mixing that we observe between pathways in the present study.

The discussion above points to the importance of being able not only to estimate the rate accurately but also the fluxes associated with pathways. In the present work, we quantify the flux through and exchange between pathways by redefining the boundary conditions when constructing the transition matrix, but history-dependent committors and reactive currents can be defined formally.^44^ It would be interesting to analyze insulin dimer dissociation within that framework. Nevertheless, the present study, along with others,^26,27,30,40,64,65^ shows that the theoretical frameworks and computational tools needed to treat complex reactions with many competing pathways quantitatively are now taking shape.

## Supporting information

Supporting_Information_rev

## Acknowledgement

The authors thank Adam Antozewski, Chatipat Lorpaiboon, John Strahan, Andrei Tokmakoff, and Jonathan Weare for useful discussions. This work was supported primarily by National Institutes of Health (NIH) award R35 GM136381. S.C.G. acknowledges support by the National Science Foundation Graduate Research Fellowship under Grant No. 2140001. This work was completed with computational resources administered by the University of Chicago Research Computing Center, including Beagle-3, a shared GPU cluster for biomolecular sciences supported by the NIH under the High-End Instrumentation (HEI) grant program award 1S10OD028655-0.

## Supporting Information Available

Extended discussion and supporting tables and figures.

