## Supporting_Information_rev for "Analysis of the dynamics of a complex, multipathway reaction: Insulin dimer dissociation"

### Supporting Information for
### Analysis of the dynamics of a complex, multipathway reaction: Insulin dimer dissociation

Kwanghoon Jeong,[†] Spencer C. Guo,[†] Sammy Allaw,[†] and Aaron R. Dinner[*,†,‡,¶]

†*Department of Chemistry, The University of Chicago, Chicago, Illinois 60637, USA*

‡*James Franck Institute, The University of Chicago, Chicago, Illinois 60637, USA*

¶*Institute for Biophysical Dynamics, The University of Chicago, Chicago, Illinois 60637, USA*

#### Rate and current estimators

To compute the reactive current, flux, and rate, we use the estimators introduced in a previous study Ref. 1 (see equations S25 and S31):

$$\begin{aligned} J^{\theta}_{AB}(s) \approx \frac{1}{2\tau|ds|} \sum_{i} \sum_{p=0}^{\frac{\tau}{\Delta}-1} & q_A(\mathbf{X}^{(i)}(\max(0, T^{-}_{D^c}(p\Delta))))q_B(\mathbf{X}^{(i)}(\min(\tau, T^{+}_{D^c}((p+1)\Delta)))) \\ & \times w(\mathbf{X}^{(i)}(0)) \left[\theta(\mathbf{X}^{(i)}((p+1)\Delta)) - \theta(\mathbf{X}^{(i)}(p\Delta))\right] \\ & \times \left[\mathbb{1}_{\{\theta\in ds\}}(\mathbf{X}^{(i)}((p+1)\Delta)) + \mathbb{1}_{\{\theta\in ds\}}(\mathbf{X}^{(i)}(p\Delta))\right]. \quad \text{(S1)} \end{aligned}$$

$$R_{AB} \approx \frac{1}{\tau} \sum_{i} \sum_{p=0}^{\frac{\tau}{\Delta}-1} q_A(\mathbf{X}^{(i)}(\max(0, T^{-}_{D^{\mathsf{c}}}(p\Delta))))q_B(\mathbf{X}^{(i)}(\min(\tau, T^{+}_{D^{\mathsf{c}}}((p+1)\Delta))))$$

$$\times\, w(\mathbf{X}^{(i)}(0)) \left[q_B(\mathbf{X}^{(i)}((p+1)\Delta)) - q_B(\mathbf{X}^{(i)}(p\Delta))\right]. \quad \text{(S2)}$$

$$k_{AB} \approx \frac{R_{AB}}{\sum_i w(\mathbf{X}^{(i)}(0)) \sum_{p=0}^{\frac{T}{\Delta}-1} q_A(\mathbf{X}^{(i)}(p\Delta))}. \quad \text{(S3)}$$

$\mathbf{X}^{(i)}$ indicates the $i$th trajectory, $\Delta$ is the saving interval (5 ps), $ds$ is a bin size of the variable $s$; we used ten bins to discretize the range of each CV, i.e. $ds = \frac{\max(\theta) - \min(\theta)}{10}$. As in previous studies,[1–3] we smooth reactive currents with a Gaussian kernel density estimate with bandwidth 0.4 $ds$ when showing projections of the currents (Figure 3 and S4). $T^{+}_{D^{\mathsf{c}}}$ and $T^{-}_{D^{\mathsf{c}}}$ are the forward and backward stopping times from time $t$, which are defined as

$$T^{+}_{D^{\mathsf{c}}}(t) = \min\{s \geq t \mid \mathbf{X}(s) \in D^{\mathsf{c}}\}$$
$$T^{-}_{D^{\mathsf{c}}}(t) = \max\{s \leq t \mid \mathbf{X}(s) \in D^{\mathsf{c}}\}.$$

#### Dependence of the results on hyperparameter choices

MSM construction involves choosing two key hyperparameters: the number of Markov states ($k$) and the lag time ($\tau$). Each of these hyperparameters involves a tradeoff between approximation and estimation errors.[4] Increasing $k$ decreases the approximation error by making the discretization finer but increases the estimation error by decreasing the amount of data associated with each state. Increasing $\tau$ decreases the approximation error by allowing more time for relaxation within a state but increases the estimation error by decreasing the number of independent time intervals within the fixed amount of data. As we describe in the main text, we go beyond a traditional MSM by incorporating memory, following Ref. 5. Increasing the number of memory terms ($\tau/\sigma - 1 = 1$, where $\sigma$ is the time interval used to compute the memory terms) decreases the approximation error but increases the estimation error as the contributions from the terms add.

As shown in Figure 4 of the main text, $k = 600$ gives estimates for the rate constant that are on par with those from $k = 2000$. To understand how the choice of $k$ and $\tau$ impacts the contributing committor estimates, we compare the estimates obtained from 100% of the data with those obtained from 50% and 75% of the data. Overall, increasing $k$ and $\tau$ decreases the consistency between the results from the full and partial datasets (i.e., increases estimation error). When $k = 200$, there is little visible difference between the estimates from 100% and 50% of the data even for the longest lag times, suggesting that the data can support a model with additional states. When $k = 600$, whether the difference is visible depends on the lag time; when $k = 2000$, the difference is visible even for the shortest lag times. To quantify the consistency, we compute the absolute value of the difference between the estimates from 100% and 50% of the data for each state and average over the 25% of states with the largest values. We see that the difference grows steadily with $k$ and $\tau$ (Figure S2, right). The values of $k = 600$ and $\tau = 2\,\text{ns}$ that we use in the main text result in a mean absolute difference of about 0.04 for the 25% of states with the largest differences.

Figure 4 in the main text shows that adding memory enables convergence of the rate constant with $k = 600$. To understand the effect of memory on the contributing committor estimates, we show how they vary with $\tau$ and the number of memory terms ($\tau/\sigma - 1$) in Figure S3. As $\tau$ and $\tau/\sigma$ increase, lower committor values are pushed closer to zero and higher committor values are pushed closer to one (which in turn makes the rate slower, as shown in Figure 4). However, at the highest values of $\tau$ and $\tau/\sigma$ considered, there is evidence that the estimation error becomes large, given that some estimates are below zero or above one. These artifacts are not visible for the choice in the main text.

Table S1: Features used to define Markov states

| **Type** | **Description** | **Residue indices**[a] **used to compute** $x$[b] | **Features**[c] |
|---|---|---|---|
| Local contacts | $\alpha$-$\alpha'$ contacts | $13-13'$<br>$09-13'$ ; $13-09'$<br>$09-09'$ | $\tanh\left(\frac{x-\mu}{2\sigma}\right)$ |
| | $\beta$-$\beta'$ contacts | $21-29'$ ; $29-21'$<br>$21-26'$ ; $26-21'$<br>$24-29'$ ; $29-24'$<br>$24-26'$ ; $26-24'$<br>$24-24'$<br>$26-26'$ | |
| | Cross contacts<br>($\alpha$-$\beta'$ ; $\beta$-$\alpha'$) | $16-29'$ ; $29-16'$<br>$16-26'$ ; $26-16'$<br>$16-21'$ ; $21-16'$<br>$09-29'$ ; $29-09'$<br>$09-26'$ ; $26-09'$<br>$09-21'$ ; $21-09'$ | |
| Global behavior | $R_{\text{com}}$ [d] | $P1-P1'$ | $x/2$ |
| | Euler angles [e] | $P1-P1'-P2'$ ; $P2-P1-P1'$<br>$P1-P1'-P2'-P3'$ ; $P3-P2-P1-P1'$<br>$P2-P1-P1'-P2'$ | $\sin(x)$, $\cos(x)$ |
| | Twisting | $26-25-25'-26'$[f]<br>$(09:11)-(12:16)-(12':16')-(09':11')$[g] | |
| | Opening[h] | $(12:16)-25-25'$<br>$25-25'-(12':16')$ | |

[a] Residue indices of chain B, where residues from different monomers are denoted with/without a prime.

[b] The raw $x$-value is a distance, angle, or dihedral depending on the number of residue indices listed (2, 3, or 4, respectively). Each residue index corresponds to the position of a $C^{\alpha}$ atom if a single index is listed. If a sequence of residues $(i:j)$ is listed, we use the geometric center of the associated backbone atoms.

[c] Functions of the raw value of a feature ($x$) used for clustering. $\mu$ and $\sigma$ are the mean and standard deviation of $x$ over the entire dataset.

[d] $P1$, $P2$, and $P3$ are defined as the center of mass of the carbonyl carbons of all residues of each monomer, A13:A19 (A-chain C-terminal $\alpha$-helix), and B9:B18 (B-chain $\alpha$-helix), respectively, following ref. 6.

[e] $P1$, $P2$, and $P3$ are defined as the centers of mass of carbonyl carbons of all residues of each monomer, A13:A19 (chain A's C-terminal $\alpha$-helix), and B9:B18 (chain B's $\alpha$-helix), respectively, following ref. 6.

[f] $\Phi_{\beta}$ in the main text.

[g] $\Phi_{\alpha}$ in the main text.

[h] $\gamma$ and $\gamma'$ in the main text.

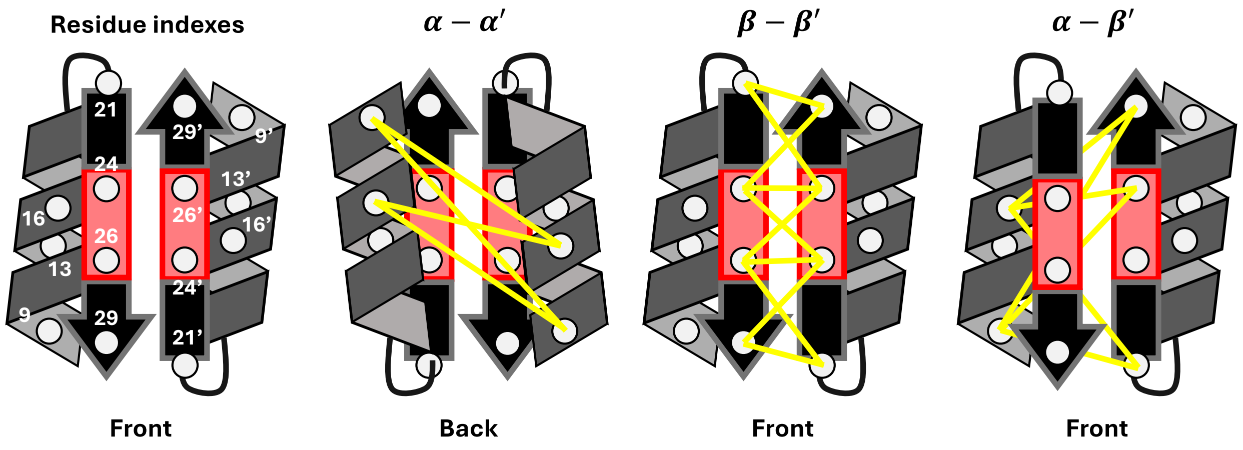


Figure S1: $C^{\alpha}$ atom pairs used to define local contact features. Only the B-chain of each monomer is shown.

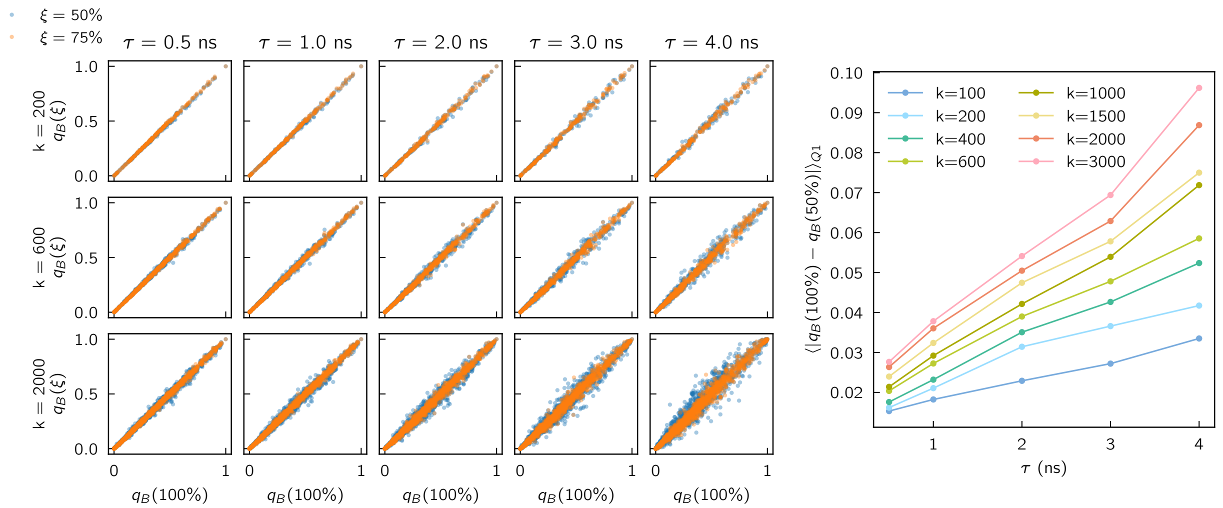


Figure S2: Committor estimates as the number of Markov states $k$ and lag time $\tau$ are varied. (left) We compare estimates obtained from all the data to those obtained from a percentage (either 50% or 75%) of the data. (right) Mean absolute value of the difference between the estimates from 100% and 50% of the data for the 25% of states (first quartile, Q1) with the largest differences.

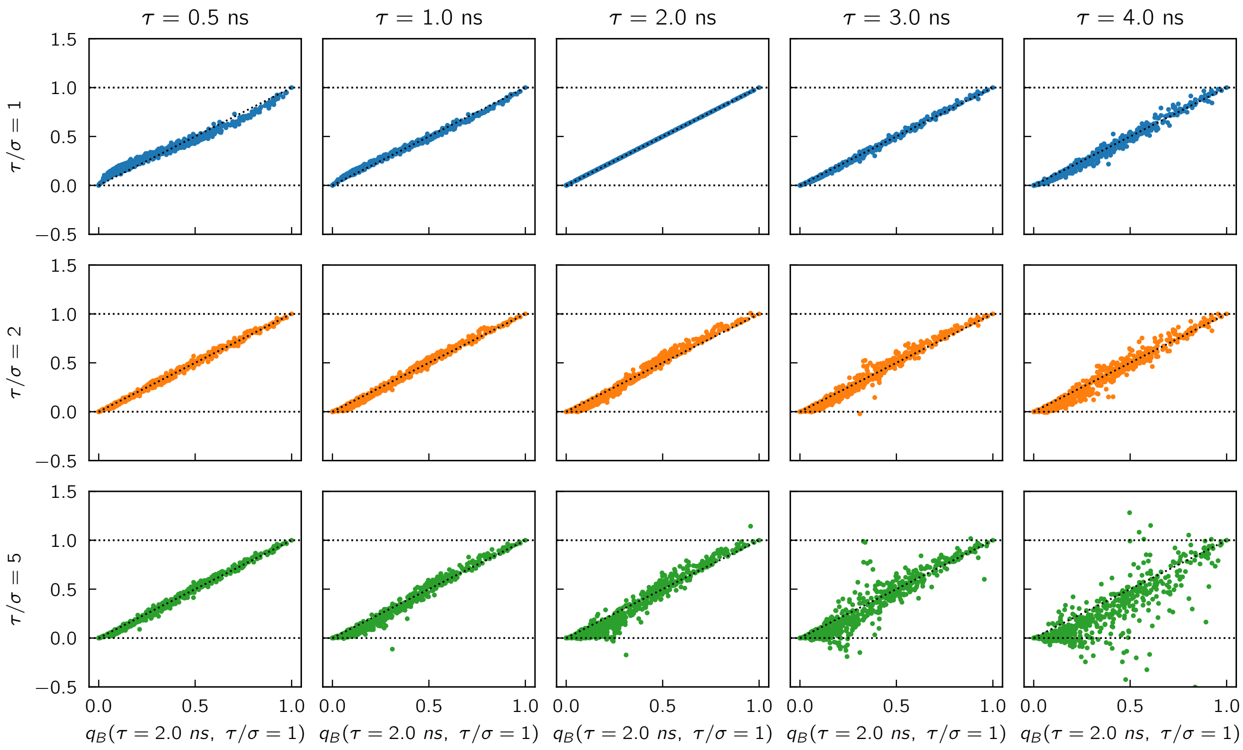


Figure S3: Committor estimates as the lag time $\tau$ and number of memory terms $\tau/\sigma - 1$ are varied. All plots are for $k = 600$.

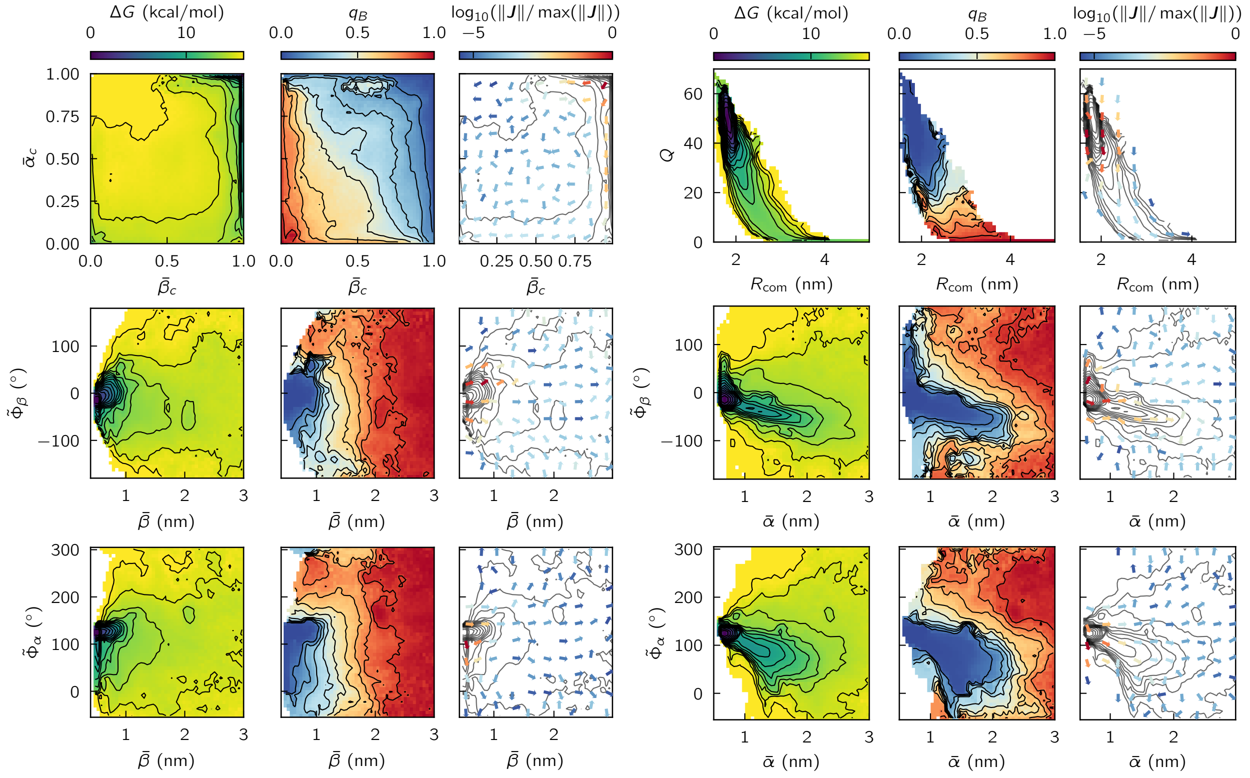


Figure S4: Projections of reactive statistics as a function of selected pairs of CVs $(x, y)$. The potential of mean force $\Delta G$ (left), committor $q_B$ (middle), and reactive current $\boldsymbol{J} = (J^x_{AB}, J^y_{AB})$ (right, arrows; contours reproduce $\Delta G$ from left) as functions of of selected pairs of CVs from previous studies.[7,8] (left and right) Contours are drawn every 1 kcal/mol. (middle) Contours are drawn every 10% increase of probability.

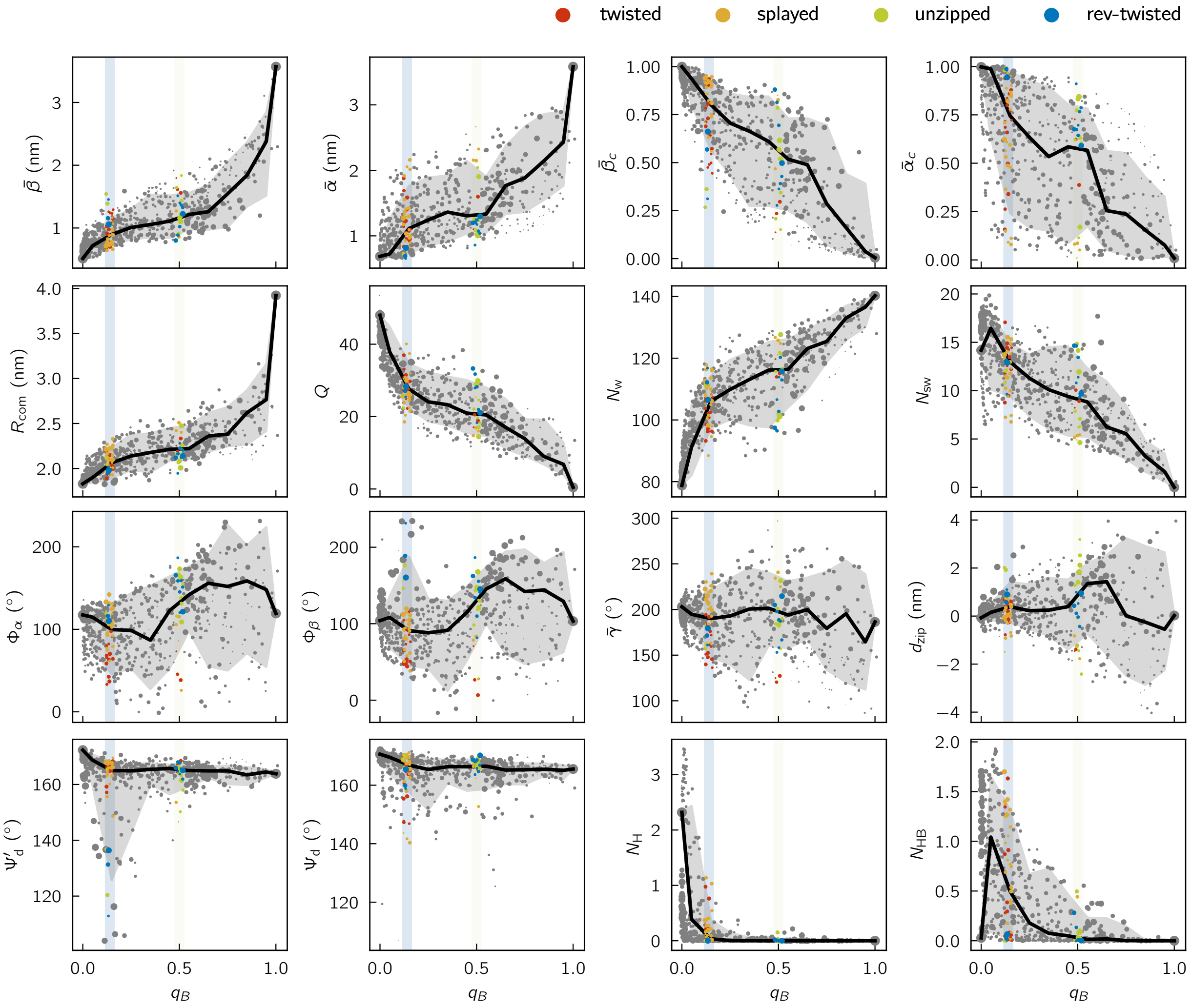


Figure S5: Evolution profiles of selected CVs. The CVs include those used to control sampling in previous studies ($\bar{\beta}$, $\bar{\alpha}$, $\bar{\beta}_{\mathrm{c}}$, $\bar{\alpha}_{\mathrm{c}}$, $R_{\mathrm{com}}$, $Q$), to describe dimer interface solvation ($N_{\mathrm{w}}$, $N_{\mathrm{sw}}$), to define the coarse-grained states ($\Phi_\alpha$, $\Phi_\beta$, $\bar{\gamma}$, $d_{\mathrm{zip}}$), to measure detachment of the $\beta$-strand and B-chain C-terminal residues ($\Psi_{\mathrm{d}}$ and $\Psi'_{\mathrm{d}}$), and to define the native hydrogen bonds ($N_{\mathrm{H}}$) and water bridges ($N_{\mathrm{HB}}$). $\Psi_{\mathrm{d}}$ is defined by the $\mathrm{C}^\alpha$ atom positions of B22, B24, and B26 (following Ref. 8), and $\Psi'_{\mathrm{d}}$ is defined correspondingly for the other monomer. A hydrogen bond is counted if its donor-acceptor distance is less than 4 Å and its donor-hydrogen-acceptor angle is greater than 150°. For $N_{\mathrm{H}}$, we consider the two backbone hydrogen bonds between B24 and B′26 and the two backbone hydrogen bonds between B′24 and B26. We define $N_{\mathrm{HB}}$ as the number of water molecules simultaneously forming hydrogen bonds between the donor and acceptor residues of the native hydrogen bonds.

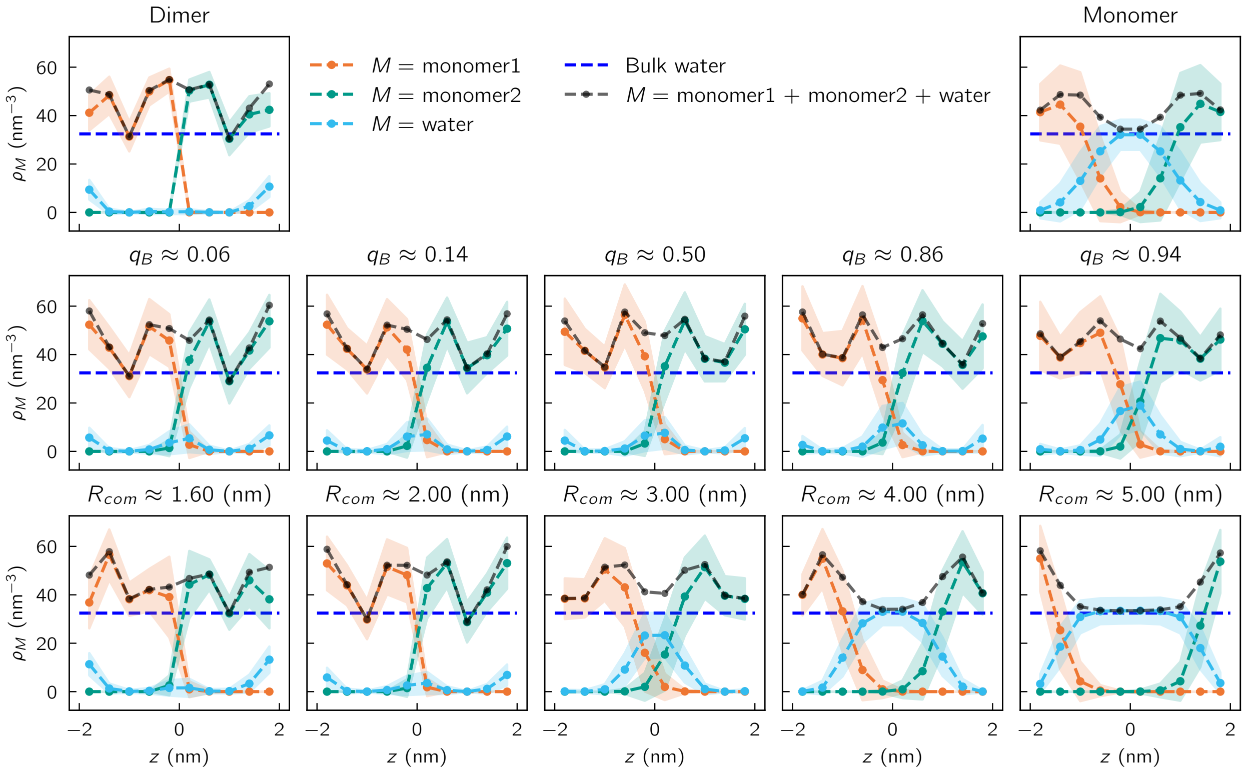


Figure S6: Number density of non-hydrogen atoms from the two insulin monomers and water molecules. The line connecting the centers of mass of the monomers is set to be $z$-axis, and the center of the two points is set to be $z = 0$. We defined a cylinder with 5 Å radius and 4 nm height so that it is centered at $z = 0$ and parallel to $z$-axis. We divide it into 10 disks and compute for each the number densities of non-hydrogen atoms from each monomer (orange and green) and the water molecules (light blue). The blue dashed line represents the bulk water density, which is taken to be the total number of water molecules divided by the simulation box volume (the volume of water excluded by the proteins is approximately 2% of the simulation box). The black dashed line shows the overall non-hydrogen atom density between the two monomers. The number densities are averaged over the structures corresponding to the state indicated above each plot. Average values are represented by the dashed lines and standard deviations are represented by shaded regions; (first row) dimer and monomer states, (second row) states corresponding to different committor values, and (third row) states with different $R_{\text{com}}$ for comparison with Ref. 9.

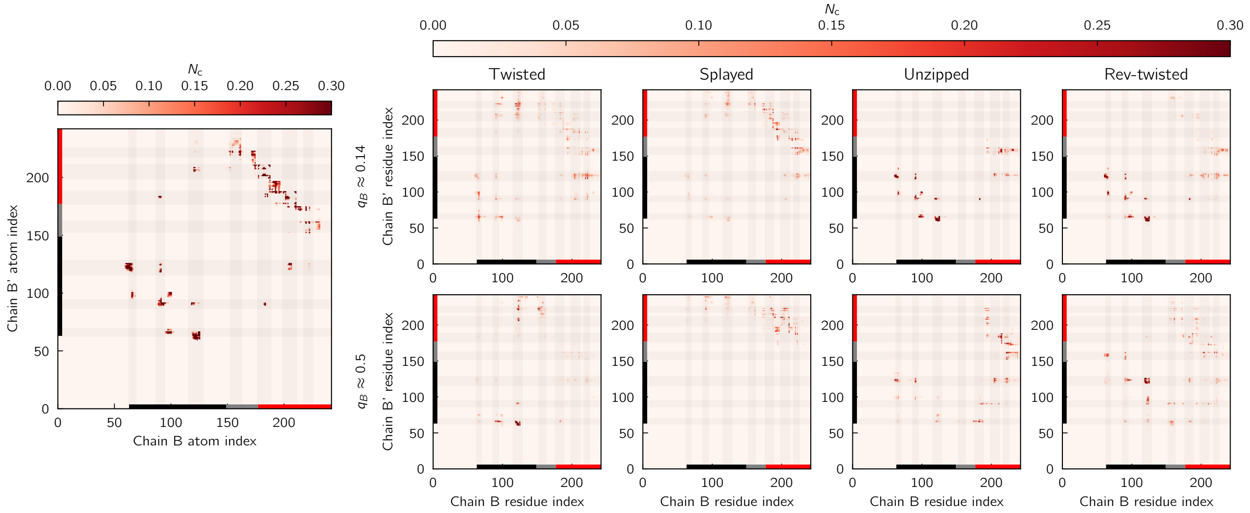

Figure S7: Atom-atom contact matrices. Shaded regions correspond to residues marked by dashed lines in Figures 6b and 8 of the main text.

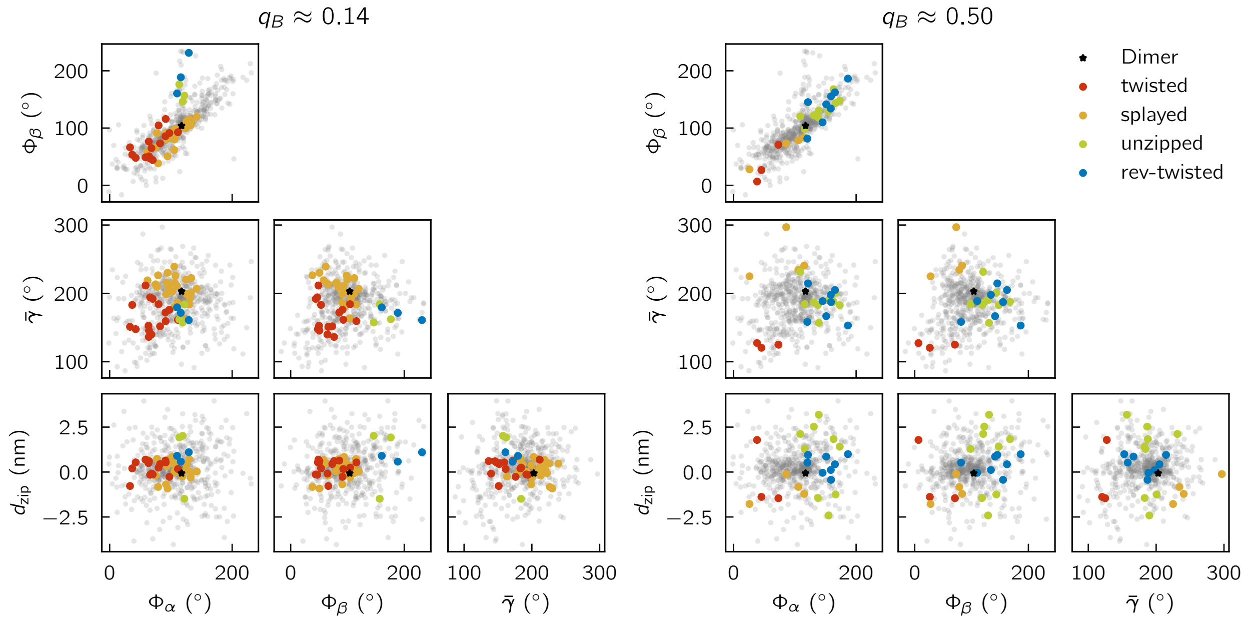

Figure S8: Projections of the Markov states (gray) in the four-dimensional space of CVs used to define the four coarse-grained states (colored). The location of the dimer state is marked with a black star. (left) $q_B \approx 0.14$. (right) $q_B \approx 0.50$.

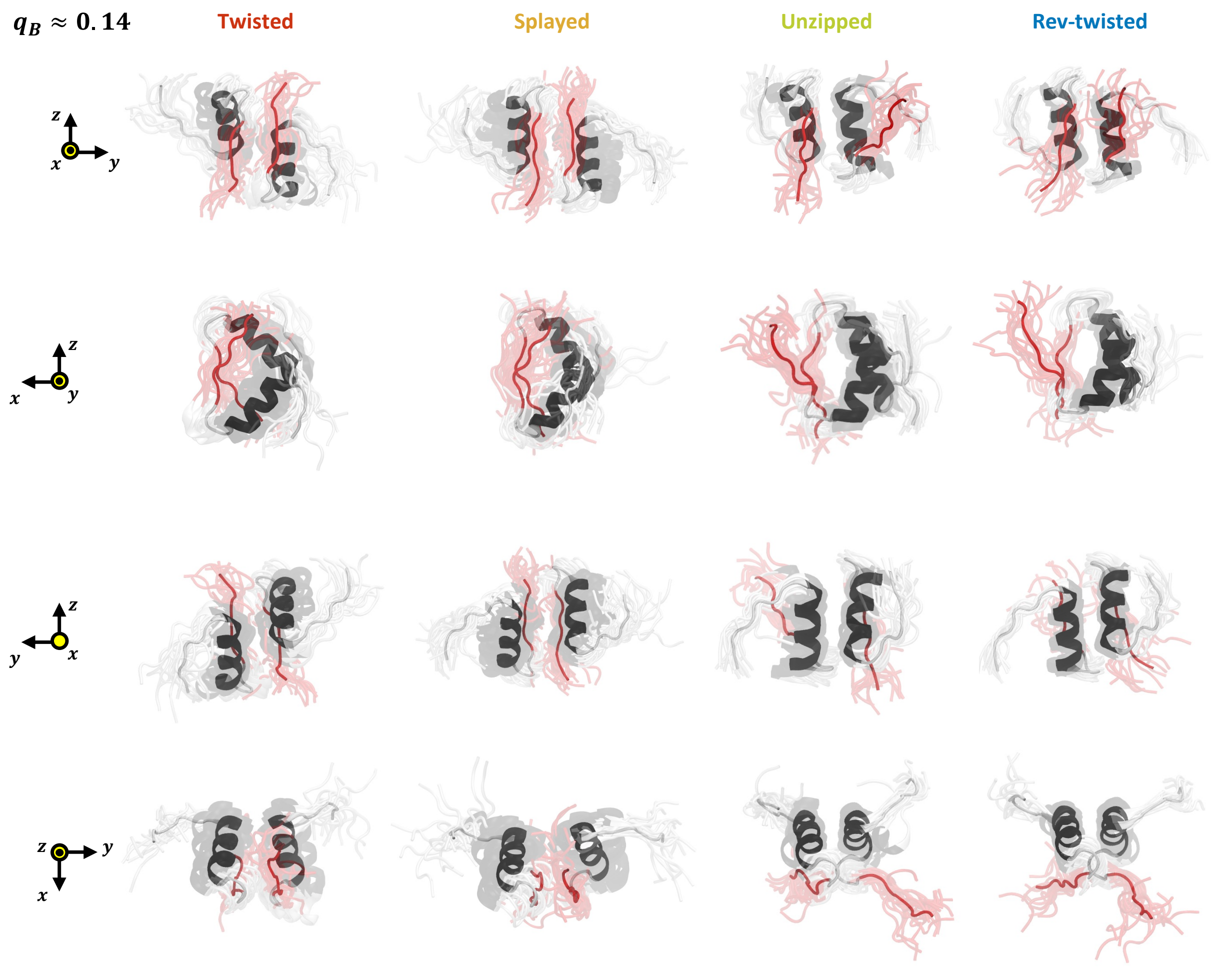


Figure S9: Different points of view of the coarse-grained states structures at $q_B \approx 0.14$.

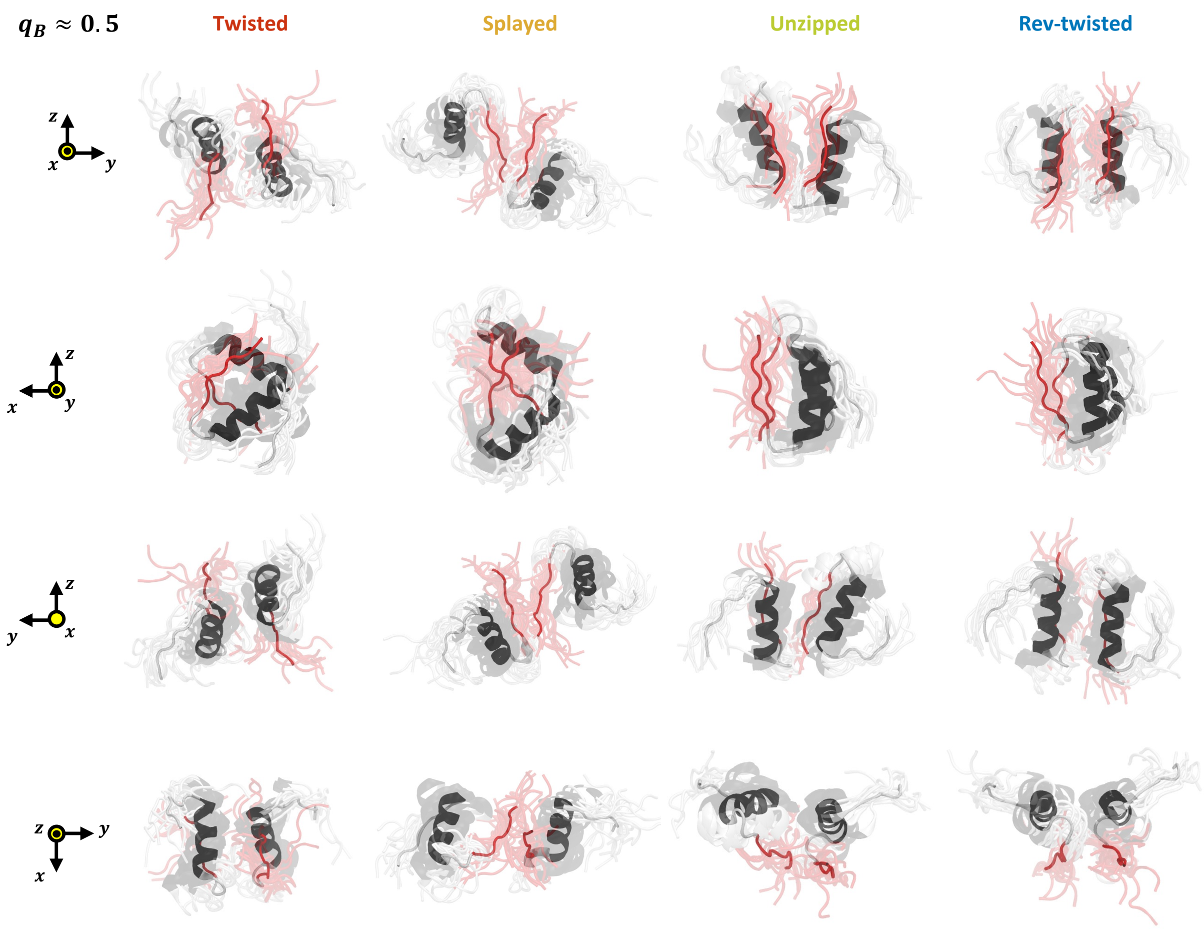


Figure S10: Different points of view of the coarse-grained state structures at $q_B \approx 0.5$.

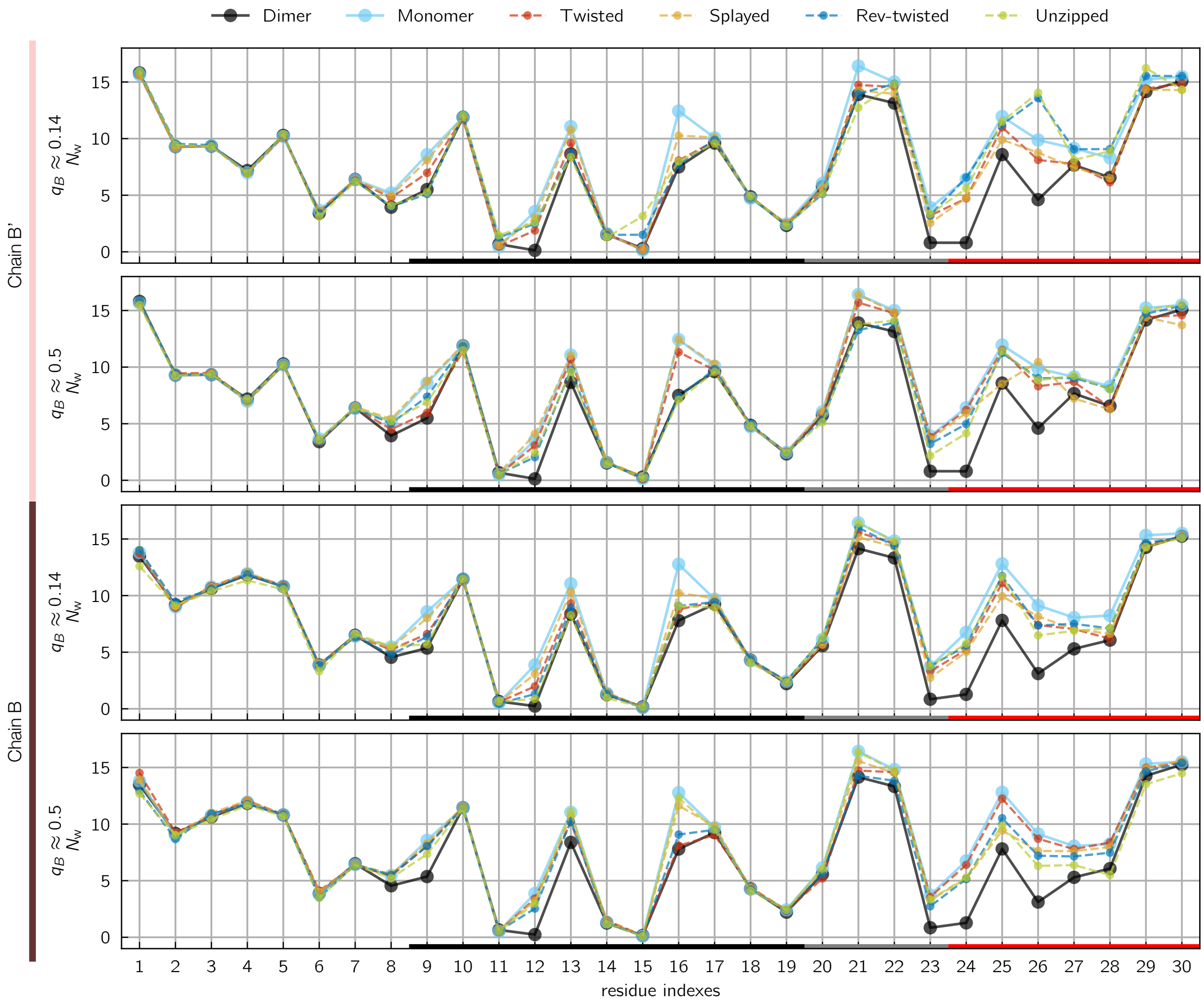


Figure S11: The number of water molecules with oxygens within 4 Å of each residue.

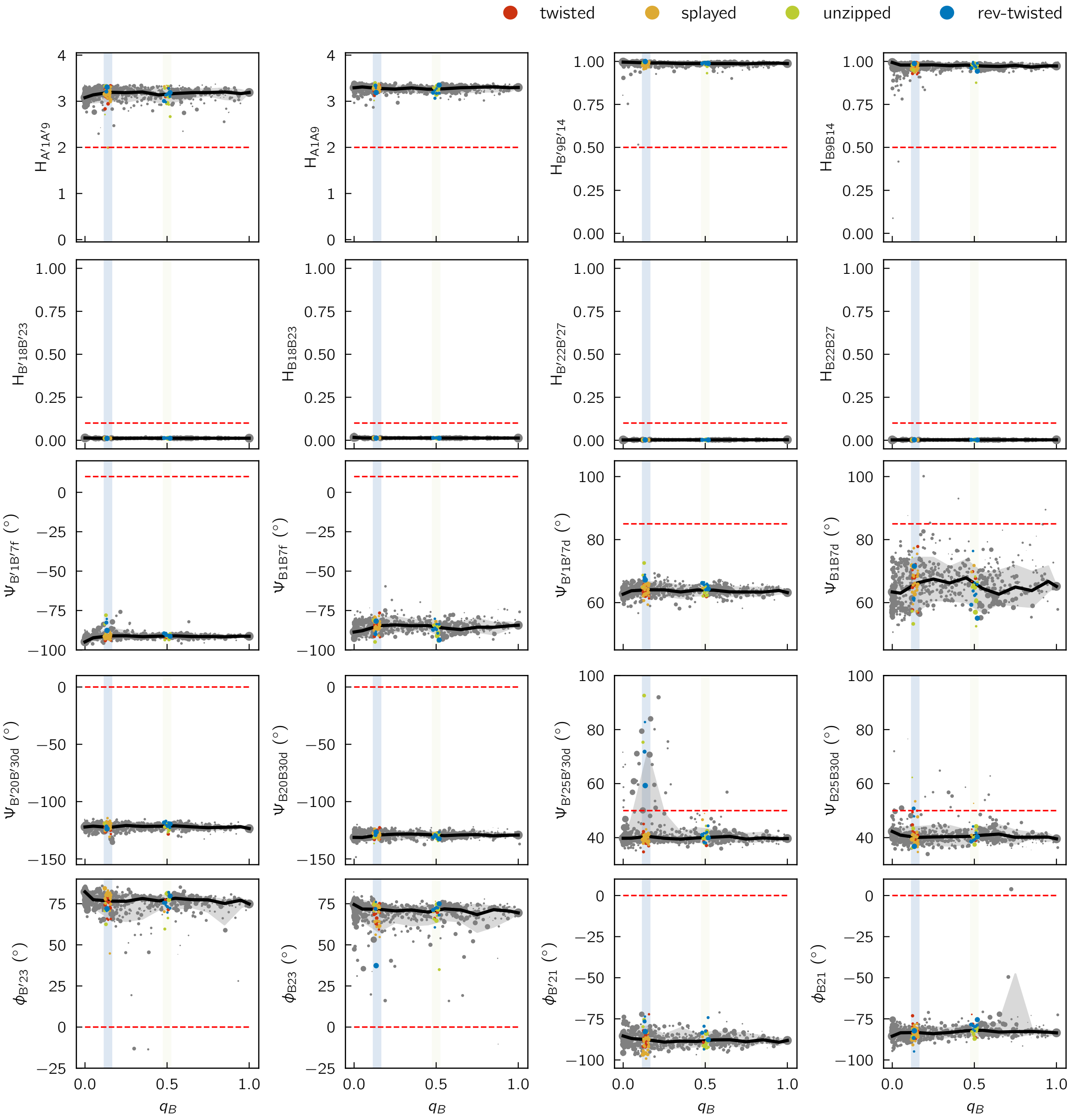


Figure S12: Evolution profiles of CVs used to describe monomer disorder in Ref. 10. Red dashed lines indicate the criteria used in that study to determine whether the indicated forms of disorder are present. (first and second rows) CVs for $\alpha$-helix melting or extension. (third row) CVs for detachment of the B-chain N-terminus. (fourth row) CVs for detachment of the B-chain C-terminus. (fourth row) CVs for the $\beta$-turn.

(8) Antoszewski, A.; Feng, C. J.; Vani, B. P.; Thiede, E. H.; Hong, L.; Weare, J.; Tok-

makoff, A.; Dinner, A. R. Insulin Dissociates by Diverse Mechanisms of Coupled Unfolding and Unbinding. *Journal of Physical Chemistry B* **2020**, *124*, 5571–5587.

(9) Banerjee, P.; Bagchi, B. Dynamical control by water at a molecular level in protein dimer association and dissociation. *Proceedings of the National Academy of Sciences* **2020**, *117*, 2302–2308.

(10) Busto-Moner, L.; Feng, C. J.; Antoszewski, A.; Tokmakoff, A.; Dinner, A. R. Structural Ensemble of the Insulin Monomer. *Biochemistry* **2021**, *60*, 3125–3136.
